# Interface swapping orchestrates carbon transfer in the archaeal acetyl-CoA decarbonylase/synthase

**DOI:** 10.64898/2026.07.07.736967

**Authors:** Erik Zimmer, Tristan Reif-Trauttmansdorff, Anthony Ciancone, Sofia Appelgren, Jörg Kahnt, Darja Deobald, Frank Abendroth, Olalla Vázquez, Georg K. A. Hochberg, Francis J. O’Reilly, Jan M. Schuller

## Abstract

The Wood–Ljungdahl pathway is one of biology’s most ancient routes for carbon fixation and energy metabolism, used by organisms such as methanogenic archaea. One of its central metabolic complexes is the acetyl-CoA decarbonylase/synthase (ACDS) complex, catalyzing acetyl-CoA synthesis and cleavage through the co-ordinated action of carbon monoxide dehydrogenase (CODH), acetyl-CoA synthase (ACS), and corrinoid iron–sulfur protein (CoFeSP). Unlike bacterial CODH/ACS, archaeal ACDS lacks a stable bifunctional CODH–ACS architecture, raising the question of how reactive CO and methyl intermediates are efficiently transferred between catalytic modules. Using cryo-electron microscopy, crosslinking mass spectrometry, small-angle X-ray scattering, and biophysical analyses, we resolved the organization and dynamics of the ∼2 MDa archaeal ACDS supercomplex from *Methanosarcina acetivorans*. We identified CoFeSP as a central architectural scaffold that self-assembles into hexa- to octameric oligomers via a conserved N-terminal region of the CdhD subunit. This scaffold likely tethers CODH and ACS through conserved disordered terminal regions, positioning the catalytic modules in the complex’s periphery. We propose a mechanism in which ACS transiently alternates between CODH and CoFeSP, enabling efficient CO and methyl-group transfer without stable binary complexes. This dynamic organization represents a fundamental difference to the stable bifunctional CODH/ACS in bacteria, highlighting how transient interactions enable efficient acetyl-CoA metabolism in archaea.

## Introduction

CO-dehydrogenase/acetyl-coenzyme A synthase (CODH/ACS) is the central enzyme complex of the reductive acetyl-coenzyme A (acetyl-CoA) pathway, also known as the Wood–Ljungdahl pathway (WLP) (1–3). CODH/ACS catalyzes the reduction of CO_2_ to CO and its condensation with a methyl-group and coenzyme A (CoA) to acetyl-CoA, thereby coupling carbon fixation with energy conservation in strictly anaerobic microorganisms, including acetogenic bacteria (acetogens) and methanogenic archaea (methanogens) (2). These organisms thrive at the thermodynamic limit of life (4, 5), often inhabiting environments that are thought to resemble early earth, where life evolved in the absence of O_2_ on inorganic gases CO_2_ and H_2_ as sources of carbon and energy, respectively (6, 7). Accordingly, the WLP is considered the most ancient CO_2_ fixation pathway, likely resembling the pathway present in life’s last universal common ancestor (LUCA) (7–9).

In the WLP, two CO_2_ molecules are reduced through distinct branches before converging at acetyl-CoA synthesis. In the methyl branch, CO_2_ is reduced to a methyl group bound to tetrahydrofolate (H_4_F) in bacteria, and tetrahydromethanopterin (H_4_MPT) or tetrahydrosarcinapterin (H_4_SPT) in archaea (10, 11). The methyl group is transferred to a corrinoid iron–sulfur protein (CoFeSP), with the assistance of a dedicated methyltransferase in bacteria, whereas archaeal CoFeSP catalyzes this transfer without an additional enzyme (10, 11). In the carbonyl branch, a second molecule of CO_2_ is reduced by ferredoxin (Fd) via a CO-dehydrogenase (CODH) to form enzyme-bound CO (1). Finally, the CO is condensed with the methyl group and coenzyme A (CoA), catalyzed by an acetyl-CoA synthase (ACS), to yield acetylCoA (1). These reactions are reversible under physiological conditions (12) (reaction 1 and Fig. 1A).

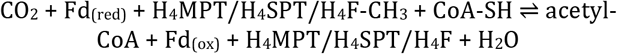

**Figure 1:**
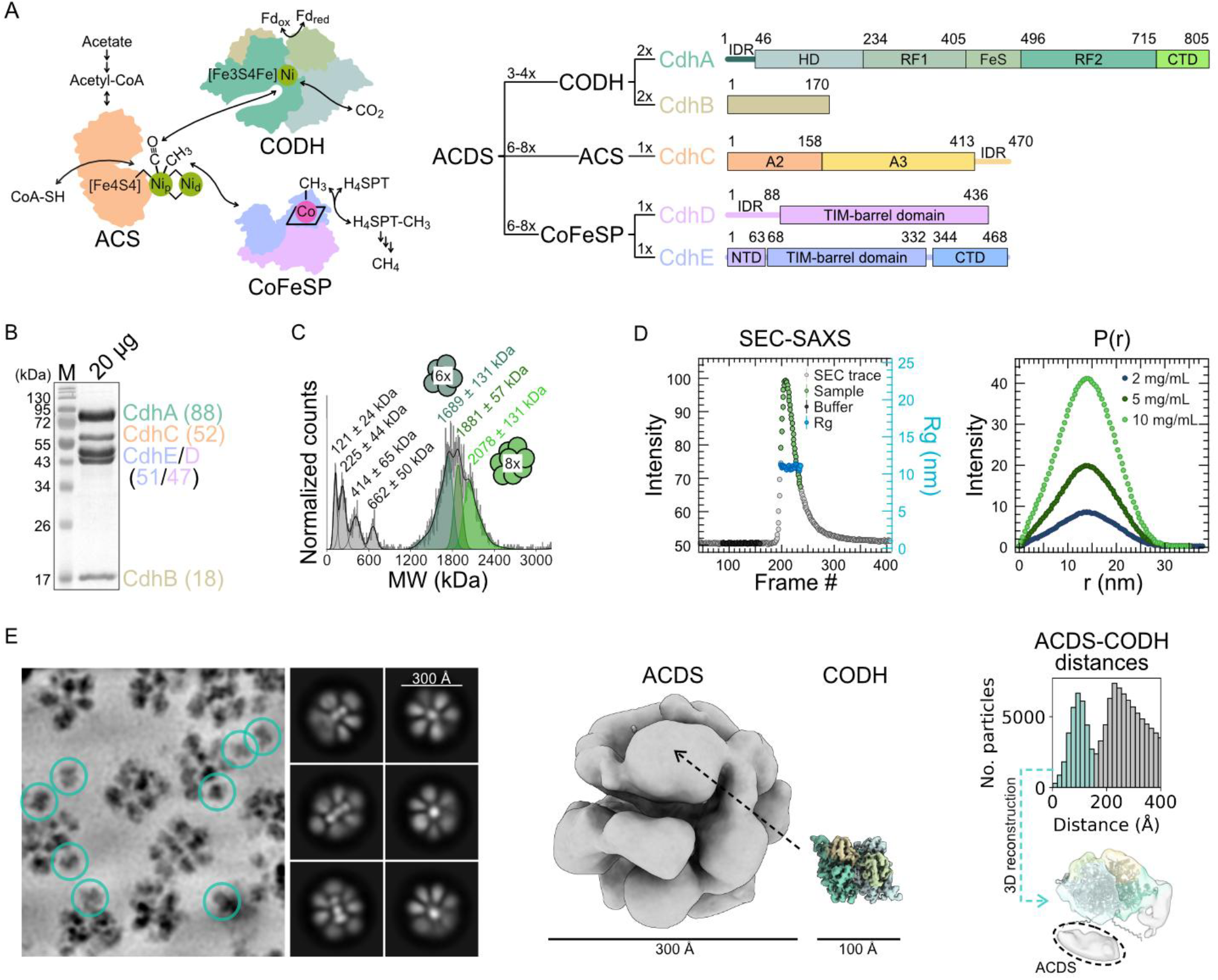
Protein biochemical and structural characterization of the M. acetivorans ACDS complex. **(A)** Partial and net reactions catalyzed by the ACDS subcomplexes ACS (yellow), CODH (shades of green), and CoFeSP (pink and blue), and the role of ACDS in aceticlastic methanogenesis. Schematic representation of the five subunits CdhA-E and their subcomplexes, domains are depicted as rectangles, unstructured regions as threads. **(B)** Coomassie-stained SDS-PAGE of ∼20 µg purified protein with protein bands labelled by their assigned subunits with their theoretical molecular weights in kDa in brackets. **(C)** Mass photometry of purified protein with fitted peaks labelled with mean ± standard deviation molecular weights. **(D)** SEC-SAXS chromatogram of total scattering intensity recorded over time with radius of gyration (Rg, blue) plotted for the protein peak, and concentration-dependent P(r) plots (distance distributions). **(E)** Representative denoised micrograph of vitrified ACDS. CODH picks are highlighted with green circles. Representative 2D classes of ACDS picks are depicted. Refined 3D reconstructions of ACDS and CODH. Distribution of distances between picks of CODH and ACDS, and 3D reconstruction of CODH particles within 150 Å of ACDS particle centers. Abbreviations: oxidized/reduced ferredoxin (Fd_ox/red_); tetrahydrosarcinapterin (H_4_SPT); coenzyme A (CoA); acetyl-CoA synthase (ACS); CO dehydrogenase (CODH); corrinoid iron sulfur protein (CoFeSP); N-terminal helical domain (HD); Rossmann-fold domain (RF); iron-sulfur domain (FeS); C-terminal domain (CTD); N-terminal domain (NTD); molecular weight marker (M); intrinsically disordered protein region (IDR).

In acetogens, acetyl-CoA supports ATP production through substrate-level phosphorylation by acetate formation (5), whereas in methanogens it functions either as an anabolic precursor or as substrate for aceticlastic methanogenesis, where it is disproportionated to CO_2_ and CH_4_ (13, 14). Notably, aceticlastic methanogenesis is restricted to the order *Methanosarcinales*, yet accounts for roughly two-thirds of the methane produced annually by methanogens (14, 15). Understanding how methanogens organize reversible acetyl-CoA synthesis and cleavage is therefore relevant to both early metabolic evolution and the modern carbon cycle.

First discovered in *Methanosarcina* spp., the archaeal CODH, ACS, and CoFeSP are organized into a ∼2 MDa multienzyme supercomplex, commonly referred to as the acetylCoA decarbonylase/synthase (ACDS) complex (16–18). This differs fundamentally from the better-characterized bacterial system, in which CODH and ACS form a stable bifunctional complex (19, 20). To date, structural insights into ACDS have been limited to isolated components and partial assemblies (21–23), leaving unresolved how this super-complex is organized and how it facilitates the efficient transfer of valuable CO and methyl intermediates between the catalytic modules.

Despite catalyzing identical reactions, the three catalytic subcomplexes (CODH, ACS and CoFeSP) differ structurally between bacteria and archaea (schematic depiction of sub-units in Fig. 1A and comparison to bacteria in Extended Data Fig. 1A). CODH holds the catalytic C-cluster, a [3Fe−4S−Ni] cluster bound to an external Fe, for reversible CO/CO_2_ conversion (19, 24–26). While bacterial CODH is a AcsA_2_ homodimer (25), archaeal CODH forms a CdhA_2_B_2_ heterotetramer with additional [4Fe–4S] clusters and domains (21). In bacteria, CODH forms a stable bifunctional complex with ACS subunit AcsB, which consists of three domains (A1-3) and houses the A-cluster, a binuclear Ni center bridged to a [4Fe–4S] cubane, where CO, a methyl group, and CoA condense to form acetyl-CoA (19, 27, 28). A protected hydrophobic tunnel running through CODH and the ACS N-terminal A1 domain connects the CODH C-cluster to the ACS A-cluster, allowing CO to pass between active sites without leakage to the surrounding (19, 20, 27). Structurally, domain A1 is a fusion of a Rossmann-fold with an alpha-helical domain called a prismane domain, which is also found in CODH subunit CdhC/AcsA (24, 29). By contrast, archaeal ACS subunit CdhC lacks the A1 domain and therefore interacts with CODH only transiently within the ACDS supercomplex (22). This raises the central mechanistic question of how archaeal ACDS maintains efficient CO transfer and catalytic coupling without the stable CODH/ACS architecture used by bacteria.

CoFeSP (corrinoid iron–sulfur protein) is a CdhDE heterodimer (AcsCD in bacteria) and provides the methyl-carrying module of the pathway, conserved both in bacteria and archaea (30, 31). CdhE consists of a C-terminal corrinoid-binding domain, a central (αβ)_8_(TIM) barrel domain, and an N-terminal [4Fe–4S] cluster domain, hypothesized to play a role in reductive reactivation of the corrinoid (30, 32, 33). Functionally, an additional methyl transferase (AcsE_2_) catalyzes the methyl transfer between the corrinoid of bacterial CoFeSP and H_4_F, whereas archaeal CoFeSP performs this reaction on its own (10, 11). Archaeal CdhD carries an N-terminal unstructured region, generally absent in bacteria (Extended Data Fig. 1B). Together with terminal unstructured regions conserved in archaeal CdhA and CdhC, this suggests that archaeal ACDS may use disordered elements to organize as a dynamic supercomplex, but their structural and mechanistic roles have remained unresolved. Here, we investigated the spatial organization of the archaeal ACDS supercomplex from the aceticlastic methanogen *Methanosarcina acetivorans* using an integrative structural biology approach. Combining cryo-EM single-particle analysis, crosslinking mass spectrometry, SAXS, and complementary biophysical techniques, we built a structural model of the ∼2 MDa supercomplex showing ACDS is organized around a CoFeSP-based oligomeric scaffold. The scaffold is formed by a coiled-coil element in the N-terminal region of CoFeSP small subunit CdhD and anchors the catalytic modules in a flexible peripheral shell. This architecture enables the ACS subunit to dynamically alternate between two mutually exclusive interaction states – a CODH-bound state positioned for CO-transfer and a CoFeSP-bound state positioned for methyl-transfer. Our findings reveal that archaeal ACDS solves the problem of intermediate channeling through scaffolded subunit dynamics, providing a distinct organizational principle for ancient acetyl-CoA metabolism.

## Results

### Native ACDS forms a flexible megadalton supercomplex

ACDS (isoform Cdh2) was natively and anaerobically purified from acetate-grown *M. acetivorans* MCD31 (34). SDS-PAGE (Fig. 1B) and tandem mass spectrometry analysis of tryptic peptides confirmed the presence of all five subunits (Extended Data Fig. 2A), while CO-dependent reduction of methylviologen (MV) (Extended Data Fig. 2B) established the activity of the CODH subcomplex. Quantification of metals with inductively coupled plasma-mass spectrometry (ICP-MS) revealed ratios of Ni, Fe, and Co that were comparable to the theoretically expected amounts for equimolar stoichiometry of all five subunits (Extended Data Fig. 2C), in accordance with previous publications (16, 17). Together, these data established that an active, metal-bound ACDS complex was purified.

Previously, the molecular weight (MW) of the ACDS complex from *Methanosarcina* species was determined by HPLC gel filtration and native PAGE (16, 17, 35). Mass photometry (MP) did not show one discrete mass, but rather several populations ranging from 1.2 to 2.4 MDa (Fig. 1C) with decreasing masses for decreasing NaCl concentrations (Extended Data Fig. 2D), similar to what had been previously reported (35). Assuming CdhA-E occur in equal stoichiometry, and CODH exists only as a CdhA_2_B_2_ dimer, the average masses observed by peak fitting approximate both a hexamer (CdhA_6_B_6_C_6_D_6_E_6_, 1540 kDa) and an octamer (CdhA_8_B_8_C_8_D_8_E_8_, 2053 kDa) of the five subunits. An octamer with one CODH dissociated (CdhA_6_B_6_C_8_D_8_E_8_, 1840 kDa) or a heptamer with 4 CODHs (CdhA_8_B_8_C_7_D_7_E_7_, 1898 kDa) fit the data as well. Populations of ACDS subcomplexes such as CODH (213 kDa) were also observed and were more abundant at lower concentrations of salt. Thus, native ACDS does not behave as a single rigid species, but as a salt-sensitive ensemble of large assemblies consistent with hexa-to oc-tameric organization. Additionally, these macromolecular properties of ACDS were shown to be unaffected by oxygen (Extended Data Fig. 2E), in contrast to the highly oxygen-sensitive catalytic activity (16).

In addition to MP, biomolecular small-angle x-ray scattering coupled to gel filtration (SEC-SAXS) was performed. From this experiment, ACDS eluted as a single peak with a homogeneous radius of gyration and an overall globular, multi-domain shape, with some degree of unstructured protein regions (Fig. 1D and Extended Data Fig. 2F). The distance distributions reported a maximum particle diameter of 306 Å, which was independent of protein concentration. A peak shoulder in the distance distributions at ∼50 Å can be explained by the presence of the three smaller subcomplexes. Collectively, our data suggest that CdhA-E assemble into oligomers of roughly 30–50 subunits, with balanced subunit stoichiometry on average. Notably, distinct oligomeric species display comparable radii of gyration.

### Cryo-EM reveals flower-shaped, flexible particles

The natively purified complex was crosslinked and vitrified for cryo-EM single-particle analysis (SPA). Particles were preferably found in thick ice and 3D reconstruction revealed a globular complex with ∼300 Å diameter (Fig. 1E and Extended Data Fig. 2), consistent with the SAXS results. This complex consists of a central density surrounded by and connected to ∼14 outer densities (Extended Data Fig. 4A). Multiple *ab initio* models and 3D classification also showed that the outer shell appears to undergo a high degree of conformational flexibility, with densities shifting relative to each other (Extended Data Fig. 4B). Refinement with applied symmetries did not improve resolution and the map could not be refined to resolutions better than 20 Å (Extended Data Fig. 4C). This led us to the conclusion that ACDS is a highly flexible supercomplex with a vast number of individual conformational states.

### CODH occurs in the periphery of the ACDS particles

Untargeted blob picking revealed early on in processing that CODH subcomplex particles could be picked individually and refined to 3.55 Å resolution when applying C2 symmetry (Extended Data Fig. 3,5A). These CODH particles were located either individually on the grid or in vicinity of ACDS particles (Fig. 1E), which is in accordance with the dissociation observed with MP (Fig. 1C). The refined CODH map revealed CODH alone, with no other subcomplexes (e.g., ACS) attached to it. *M. acetivorans* CODH closely compares to the previously reported structures of archaea-type CODH from *M. thermophila* (22), *M. barkeri* (21), *Candidatus* Ethanoperedens thermophilum (23), and *A. fulgidus* (36), with Cα superposition root mean square deviations (rmsds) of 1.1-1.7 Å (Extended Data Fig. 5B). CODH occurs as a CdhA_2_B_2_ heterotetramer with a diameter of ∼100 Å. Furthermore, the cryo-EM map features densities confirming the presence of all expected [4Fe-4S] B-, D-, E- and F-clusters and the catalytic Fe-[Ni-3Fe-4S] C-cluster.

To analyze the CODH particles that are associated with ACDS, we plotted the distances between CODH and ACDS center-to-center particle picks within each micrograph. This revealed a population of CODH with an average CODH-ACDS distance of ∼100 Å (Fig. 1E), which is in accordance with the distance from ACDS particle centers to the centers of the outer sphere densities. Refinement of the CODH map with only the particle population in vicinity to ACDS showed that CODH appears to be oriented with its N-terminal disordered region pointing towards the center of the ACDS particles (Fig. 1E, Extended Data Fig. 3).

### Crosslinking mass spectrometry captures ACS in the methyl- and CO-transfer states

Since the low-resolution SPA map did not allow ACDS model building, we used photo-crosslinking mass spectrometry (37) (crosslinking MS) under strictly anaerobic conditions to define interactions between the three distinct catalytic building blocks: CODH, ACS, and CoFeSP. We identified 855 crosslinks within the complex (1 % residue-pair FDR). These comprise 763 homomeric, non-overlapping, 23 homomeric overlapping, and 78 heteromeric (inter-subunit) crosslinks (Extended Data Fig. 6A, and Fig. 2A). The homomeric crosslinks were in accordance with both predicted intra-subunit distances derived from AlphaFold2 (38, 39) models and in structures of bacterial homologs (Extended Data Fig. 6A).

Among the crosslinks between identical peptide sequences (indicating homomeric contacts), 17 were found in ACS subunit CdhC. The majority of these crosslinks were clustered in ACS domain A3 housing the A-cluster, consistent with homo-oligomerization or crowding within the ACDS complex (Extended Data Fig. 6C). Three such cross-links to identical sequences were also identified in CoFeSP subunit CdhE (see following section, Extended Data Fig. 6D). More informatively, the heteromeric crosslinks were dominated by contacts between ACS and either CODH or CoFeSP. The majority of heteromeric crosslinks were clustered between CdhC and either the CODH large subunit, CdhA, or the CoFeSP small subunit, CdhD. This indicated that the cross-linking captured two states in the enzyme’s catalytic cycle: The state of methyl-transfer from the A-cluster to CoFeSP, and the state of CO-transfer from the A-cluster to the C-cluster, harbored in CODH. We therefore used AlphaLink2 (40) (AL2) to model these two distinct ACS interaction states.

AL2 generated a confident ACS-CODH model (pLDDT 0.89, pTM 0.85), with 11 of the CdhA-CdhC crosslinks satisfying the 25 Å Cα-Cα distance restraint of the crosslinker (37, 41) (Fig. 2B). The model is in accordance with the cryo-EM structure of CODH-ACS from *M. thermophila* (22) (CdhA-C iRMSD 2.989 Å (42)) (Extended Data Fig. 7A). Just like in the cryo-EM structure, the interaction surface in the AL2 model is formed mainly between the methanogen-specific CdhA C-terminal domain (21), and CdhC domain A2, as well as between CdhA Rossmann-fold domain 2 and CdhC domain A3 (Fig. 2B). This arrangement positions the A-cluster next to the exit site of the hydrophobic tunnel within CODH, establishing a connection to the C-cluster for CO-transfer. Compared to bacterial bifunctional CODH/ACS, the modeled ACS conformation resembles that of the closed conformation of CODH/ACS (Extended Data Fig. 7C), the conformation in which CO-transfer/carbonylation of the A-cluster occurs (22, 27, 43). The position and function of bacterial ACS prismane domain A1 is taken over by CdhA Rossmann-fold domain 2 in archaea. Thus, despite lacking the A1 domain, archaeal ACDS appears to generate a functionally analogous CO-transfer state through a distinct CODH–ACS interface. Interestingly, one highly confident CdhA-C crosslink was found between CdhA residue E15 and CdhC residue K452, both located in terminal disordered regions, suggesting that flexible extensions may help position or tether ACS during this state (Fig. 2D).

**Figure 2:**
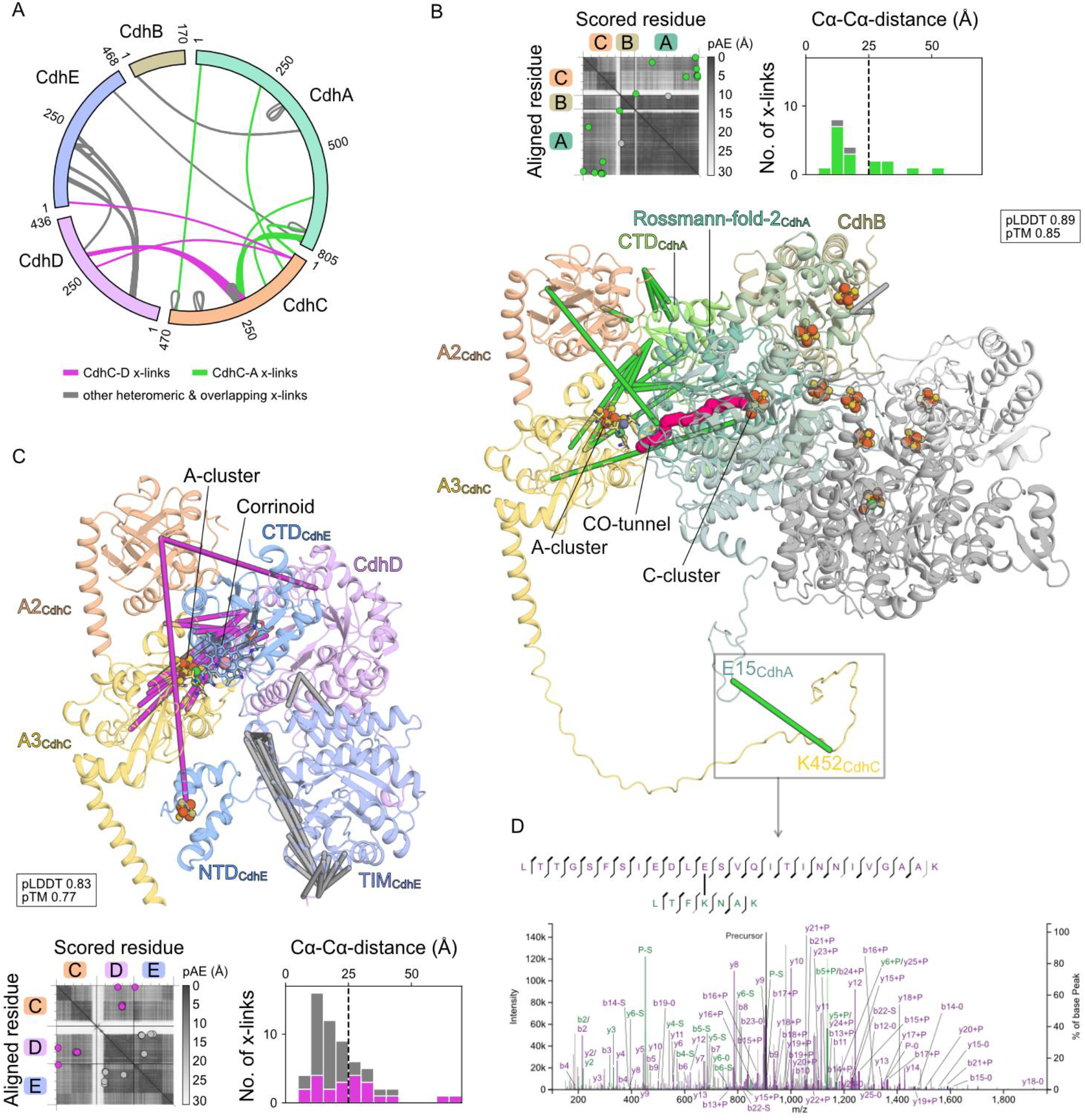
Crosslinking mass spectrometry (crosslinking MS) of the purified ACDS complex. **(A)** Circle plot depicting all self-overlapping and heteromeric crosslinks found for the five ACDS subunits (CdhA-E) at a 1 % residue-pair level false-discovery rate. **(B)** AlphaLink2 top models of CdhC (shades of yellow) in complex with CdhA_2_B_2_ (shades of green, second protomer in grey), **(C)** or in complex with CdhDE (pink and blue). Cofactors were created from superpositions of related structures (pdb-ids 1oao and 3cf4 for A- and C-clusters, 2ycl for CdhE corrinoid and [4Fe-4S]) on respective domains, the CO tunnel was predicted with Caver 2.0. pAE-plots and crosslink distance distributions are shown above or below the models, with vertical dashed lines indicating the expected crosslink threshold of 25 Å. Crosslinks are depicted as solid lines, CdhC-D crosslinks in purple, CdhC-A crosslinks in green. **(D)** Sequence and peptide-spectrum match of the crosslink identified between E15 and K452 of the terminal disordered regions of subunits CdhA (green) and CdhC (yellow) shown in **(B)**. Abbreviations: C-terminal domain (CTD); N-terminal domain (NTD); TIM-barrel domain (TIM); predicted aligned error (pAE).

A separate AL2 model of ACS bound to CoFeSP was also well supported (pLDDT 0.83, pTM 0.77), with 11 of the CdhC-CdhD crosslinks satisfying the 25 Å Cα-Cα distance restraint (Fig. 2C). In this model, ACS engages CoFeSP primarily through contacts between the (αβ)_8_-barrel of CdhD and ACS A3 domain, where all observed CdhC-CdhD crosslinks are located. The N-terminal [4Fe-4S] domain of CdhE is also predicted to contribute to the interaction. When comparing our model of CoFeSP with the transfer-competent resting state (33) we see a rotation and shift in the B12-binding C-terminal domain (CTD) of CdhE, revealing a similar flexibility compared to its interaction with either the methyl-donating methyltransferase (32) or the reductive activator RACo (44), although there are no heteromeric crosslinks in the CTD (Extended Data Fig. 7D). Superposition of the different domains in the AL2 model with experimental structures of bacterial ACS and CoFeSP (21, 27, 33) positions Co in the corrinoid in proximity to the A-cluster. The predicted CdhC conformation in this model resembles that of the open conformation found in bacterial CODH/ACS (Extended Data Fig. 7C), the conformation in which (de)methylation of the A-cluster occurs (45). Recently, the structure of *Clostridium autoethanogenum* CoFeSP in complex with CODH/ACS has been determined (43), which closely resembles the model obtained for *M. acetivorans* with the CdhD and CdhE N-terminal domain establishing the interaction (CdhC-D iRMSD of 1.698 Å (42)) (Extended Data Fig. 7B). In the bacterial structure, a third interface is formed between CdhC domain A1 and another face of the CdhD (αβ)_8_-barrel (43), which is lacking in the methanogenic model due to the absence of domain A1. This points towards a higher affinity of acetogenic ACS to CoFeSP compared to methanogens, which is perhaps necessary since both proteins are not permanently connected in one complex.

To test whether ACS could engage CODH and CoFeSP simultaneously, we also generated an AL2 model using all heteromeric crosslinks, which predicted ACS in an extended conformation (pLDDT 0.82, pTM 0.62, Extended Data Fig. 6B). In this model, CODH is only in contact with CdhC domain A2, and CoFeSP is only in contact with CdhC domain A3, hence both interaction interfaces are smaller compared to the separate models. This model fulfills only 5 of the CdhC-CdhA and 7 of the CdhC-CdhD crosslinks (< 25 Å Cα-Cα distance), and superposition reveals that the A-cluster is positioned neither in proximity to the corrinoid nor to the CO-tunnel. We therefore conclude that the ACS–CODH and ACS–CoFeSP assemblies represent mutually exclusive catalytic states. For CO-transfer, ACS must disengage from CoFeSP and bind CODH; for methyl-transfer, ACS must disengage from CODH and bind CoFeSP. This alternating mechanism contrasts with bacterial CODH/ACS, in which constitutive CODH-ACS interaction is maintained by ACS domain A1 (20, 27). Since all ACDS subcomplexes are co-purified, the intact supercomplex must be stabilized by additional uncharacterized structural elements.

### CoFeSP oligomerizes through a coiled-coil formed by the N-terminal disordered region of subunit CdhD

Previously, it was shown that the methanogenic ACDS could be resolved into its three functional subcomplexes: CODH, ACS and CoFeSP, either using detergents (46, 47) or by limited proteolysis (11). The detergent resolution protocol was reproduced to resolve the three subcomplexes, as evidenced by SDS-PAGE (Fig. 3A). After subsequent buffer exchange for detergent removal, the proteins were characterized by mass photometry (Fig. 3A). CODH and ACS appeared as the expected heterotetramer of ∼210 kDa and monomer of ∼50 kDa, respectively. In contrast, an oligomerization pattern of CoFeSP was observed, with distinct populations fitting a CoFeSP monomer, dimer, trimer, and tetramer (98-392 kDa). To the best of our knowledge, no oligomerization of bacterial CoFeSP has been reported.

**Figure 3:**
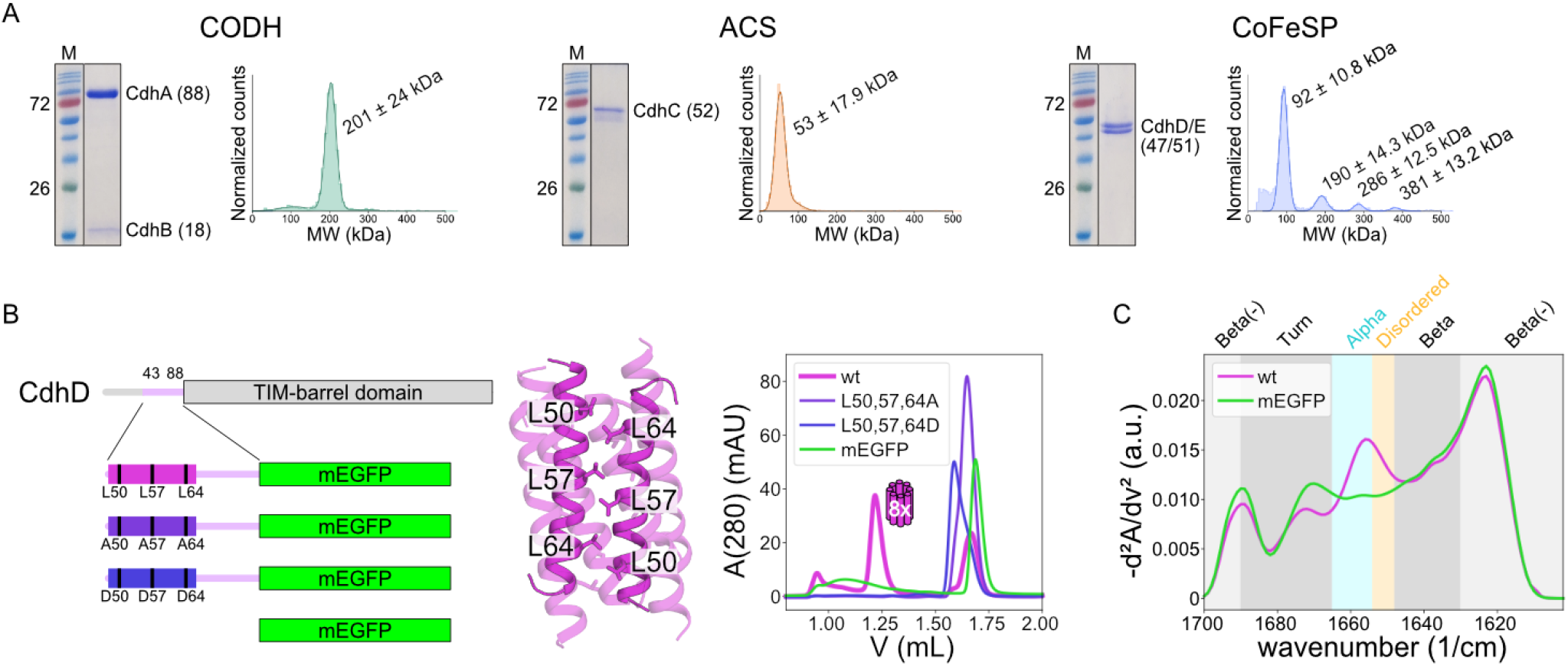
Characterisation of resolved subcomplexes and the CdhD N-terminal disordered region. **(A)** Coomassie-stained SDS-PAGE gels and mass photometry histograms of detergent-resolved subcomplexes CODH, ACS, and CoFeSP. Fitted mass photometry peaks are labelled with their mean ± standard deviation molecular weights. **(B)** Schematic representation of the purified CdhD-mEGFP fusion constructs and AlphaFold2 prediction of a coiled coil formed by CdhD residues 44-70, with mutated Leu residues labelled. Superdex 200 Increase 3.2/300 gel filtration chromatograms of purified CdhD N-terminus fusion construct with mEGFP (wt, pink), L to A (violet) and L to D (blue) mutants, and mEGFP alone (green). **(C)** MMS (microfluidic modulation spectroscopy) second derivative spectra of mEGFP (green) and CdhD N-terminus tagged mEGFP (wt, pink). Highlighted wavenumber ranges correspond to the vibrations of certain secondary structural elements. Abbreviations: acetyl-CoA synthase (ACS); CO dehydrogenase (CODH); corrinoid iron sulphur protein (CoFeSP); wildtype CdhD_IDR_-mEGFP (wt); mutated CdhD_IDR_-mEGFP (L50,57,64A, L50,57,64AD).

When comparing AlphaFold2 predictions of representative archaeal with bacterial CoFeSP small subunits, we found that the N-terminal disordered region is conserved in archaeal CdhD, but not in bacterial AcsD (Extended Data Fig. 1B). This led us to hypothesize that this region could be responsible for oligomerization. Indeed, AlphaFold2 predictions of the CdhA, CdhC, and CdhD conserved terminal disordered regions resulted in a 14-residue α-helix of CdhD residues 45-68 in the otherwise mostly disordered regions predicted to form a hexameric or octameric antiparallel coiled coil with medium confidence (Extended Data Fig. 8A). The predicted coiled coil is formed by a Leu-zipper (Leu residues 50, 57, 64) and other hydrophobic residues.

To test the existence of an oligomerization site, fusion constructs of part of the CdhD N-terminal disordered region (residues 43-88, CdhD_IDR_) with monomeric enhanced GFP (mEGFP) (48) were purified from *E. coli* (Fig. 3B and Extended Data Fig. 8B,D). While mEGFP alone eluted as the expected monomer in gel filtration, CdhD_IDR_-mEGFP showed an additional, distinct peak at higher apparent molecular weight (Fig. 3B). Gel filtration calibration indicated a MW of 274 kDa, while mass photometry of that peak indicated a MW of 215 kDa (Extended Data Fig. 8C), fitting octameric (272 kDa) and hexameric (204 kDa) oligomers, respectively. When Leu residues 50, 57 and 64 were mutated to either Ala or Asp, oligomerization was lost. In addition, increased α-helical and disordered contents were observed for CdhD_IDR_-mEGFP compared to mEGFP with microfluidic modulation spectroscopy (MMS) (49) (Fig. 3C). Taken together, the conducted experiments indicated the presence of the predicted hexa- or octameric coiled coil formed by CdhD residues 45-68, creating an oligomerization interface with stoichiometry in accordance with that observed for the whole ACDS complex.

### Integrative modelling visualizes the putative mechanism of efficient intermediate channeling in ACDS

To combine the experimental results into models of potential ACDS conformations, integrative models of an ACDS octamer (CdhA_8_B_8_C_8_D_8_E_8_) were simulated using Assembline (50) (workflow shown in Extended Data Fig. 9A). Although the modelling converged, sampling accuracy was low due to the low confidence fitting of rigid bodies into the cryo-EM map (Extended Data Fig. 9B,C); however, the top model was in close agreement with experimental crosslinking MS and SAXS data (Fig. 4B). Deviations likely arose from the comparison of a single modelled conformation to the continuum of conformations and oligomeric states of the ACDS super-complex observed experimentally. Despite the low sampling accuracy, the CdhD coiled coil was consistently positioned in the cryo-EM map center, due to optimization of connectivity restraints to the CoFeSP structured domains which, together with the other rigid bodies, were found in the outer shell of the complex (Fig. 4A).

**Figure 4:**
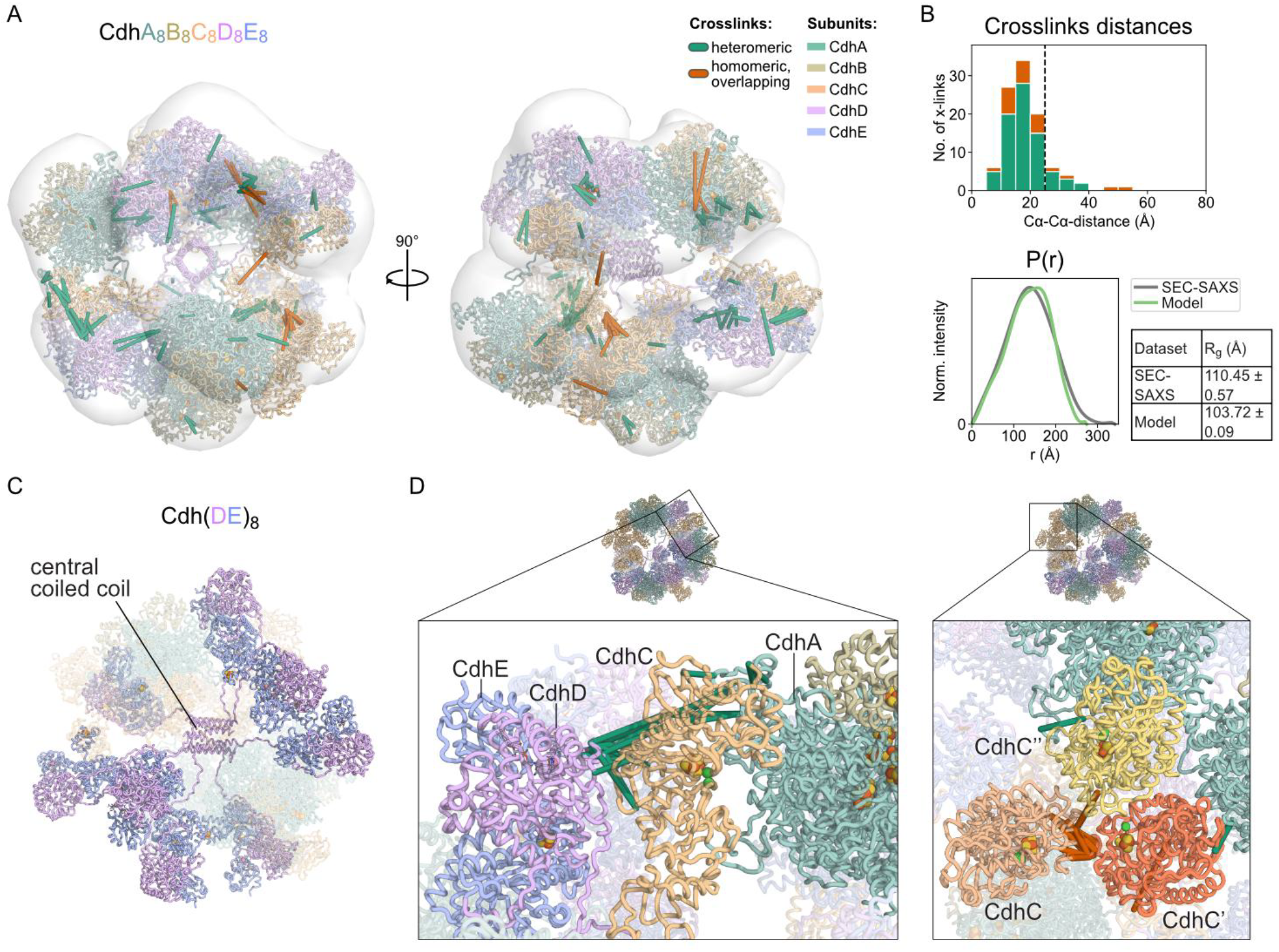
Integrative modeling of the 8-fold ACDS supercomplex. **(A)** Overview of the protein model, depicted as cartoon tubes, colored by subunit (CdhA green, CdhB olive, CdhC yellow, CdhD pink, CdhE blue). The Gaussian-filtered cryo-EM map is depicted as transparent surface, all 101 heteromeric (green) and homomeric overlapping (orange) crosslinks are depicted as dashes between their minimal-distance residue pairs. **(B)** Crosslink distance distribution, with the vertical dashed line indicating the expected crosslink threshold of 25 Å, and *P(r)* distance distribution simulated from the supercomplex model including superimposed cofactors, compared to the experimentally obtained *P(r)* plot from SEC-SAXS. Experimental and simulated radii of gyration (R_g_) are presented. **(C)** Cartoon model of the scaffold formed by the CoFeSP small subunit CdhD (pink) with its centrally positioned coiled coil. Subunits CdhA-C are shown with transparency. **(D)** Closeups of modelled CoFeSP-ACS-CODH (right) and ACS trimer (left) interactions and depiction of supporting crosslinks. Cofactors were created from superpositions of related structures (pdb-id 9c0s for A- and C-clusters, 2ycl for CdhE corrinoid and [4Fe-4S]) on respective domains.

The modelling provided potential conformational states of the ACDS complex, in which the oligomer is held together by anchoring of subunit CdhD, through its coiled coil, to the center of the complex (Fig. 4C). This allows the catalytic components to maintain a high degree of conformational flexibility while simultaneously remaining bound together and in proximity, exemplified by different subcomplex interactions found in the outer shell of the complex (Fig. 4D). Although there is a lack of protein biochemical data on the disordered regions of CdhA (residues 1-43) and CdhC (residues 401-470), we propose their interaction with the CdhD disordered protein region prevent dissociation of CODH and ACS. All disordered regions reside within a mutually accessible interaction sphere (Extended Data Fig. 9D). This organization implies that the complex is not only stabilized but also configured to support efficient channelling of reactive intermediates between the catalytic modules.

To test whether this architectural arrangement indeed translates into functional coupling during catalysis, we measured CO leakage from the ACDS complex during acetylCoA synthesis by addition of hemoglobin, as previously described for the bifunctional CODH/ACS complex (51). We found that CO-hemoglobin formation was necessitated by the decrease of acetyl-CoA formation activity, and that the rate of acetyl-CoA formation was not affected by the addition of hemoglobin (Extended Data Fig. 9E). This suggests that CO only escapes from the complex when ACS is no longer capable of acetyl-CoA synthesis due to limitation of the provided methyl-cobinamide. Taken together, the mode of catalysis employed by the ACDS complex appears to be as efficient in catalytic intermediate transfer as the bifunctional CODH/ACS.

### Phylogenetic analysis reveals the broad distribution of a multitude of CODH/ACS/ACDS variants across both domains of prokaryotes

To analyze the distribution of the two types of CODH/ACS and ACDS systems across archaea and bacteria, we conducted a phylogenetic analysis of acetyl-CoA synthases (Fig. 5A). Despite the historical classification of the two-domain (A2-3) CdhC as archaeal and the three-domain (A1-3) AcsB as bacterial (29, 52), we conclude that both enzymes occur in both the bacterial and archaeal domains. AcsB was found in archaeal Bathyarchaeota (TACK), Methanomicrobia, Methanomassilicoccales, Thermoplasmatota, Halobacteria (all Euryarchaeota), and Heimdalarchaea (Asgard), and bacterial CdhC was found in Chloroflexota, Thermodesulfobacteriota and Myxococcota. Interestingly, we identified a clade of CdhC fused to a prismane domain, thus resembling the bacterial AcsB; however, this CdhC clusters with other CdhCs and holds the CdhC C-terminal extension, while the prismane domain separately clusters with AcsB-type prismane domains (Fig. 5B). Depending on where the root of the phylogeny lies, the prismane domain has either been gained twice (root in CdhC) or represents the ancestral state (root in AcsB), which was lost and then regained. The prismane-CdhC is found in gene clusters holding two CdhC, one with and one without the prismane domain, and an archaeal type CODH.

**Figure 5:**
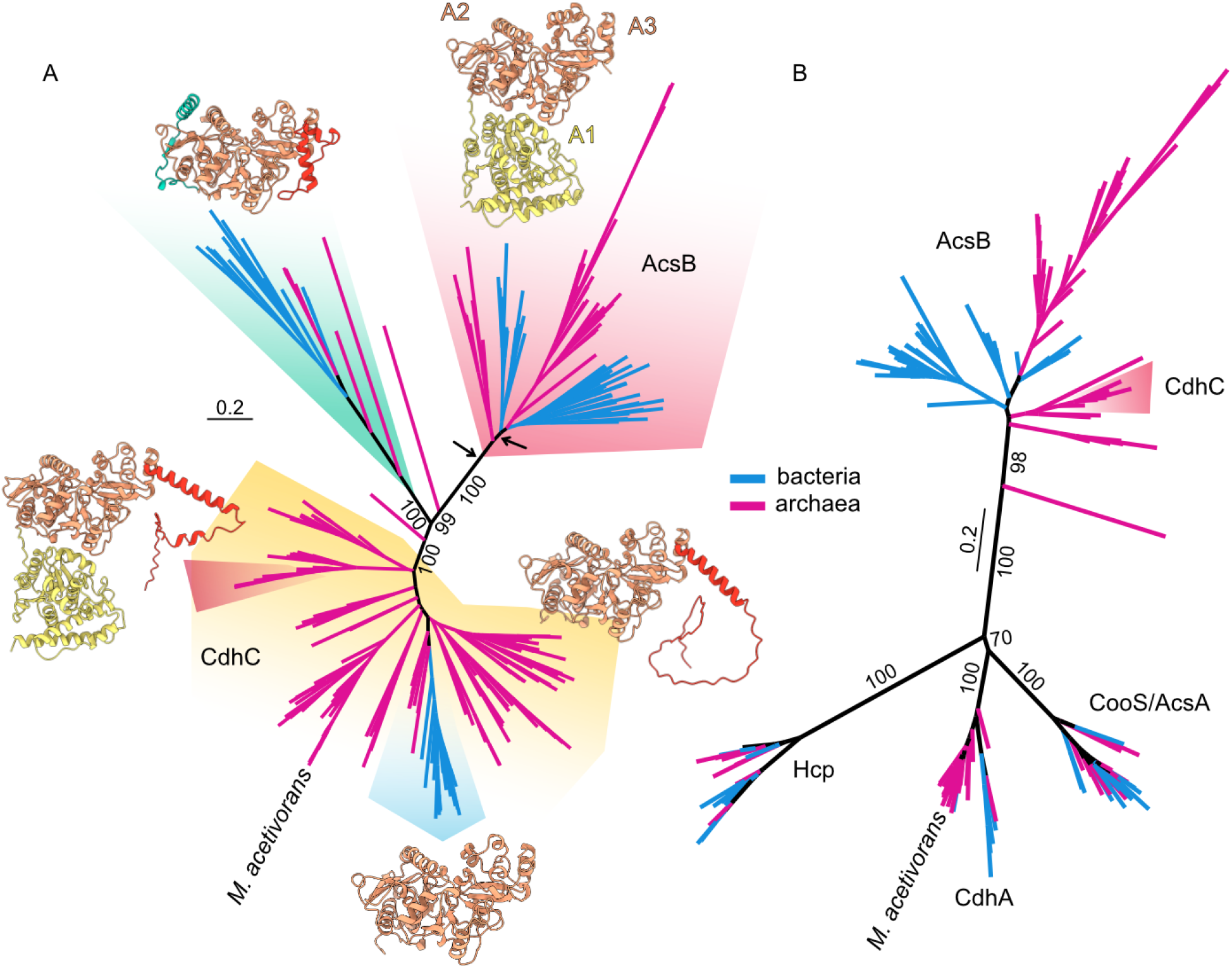
Structural phylogenies of ACS and prismane domains. **(A)** Phylogenetic tree of ACS. Blue and pink branches lead to proteins found in bacterial and archaeal genomes, respectively. Arrows point to proposed roots. Two clades are ACS core domains A2 and A3 (orange) N-terminally fused to prismane domains (yellow), and these are indicated by pink triangles. One clade holds an N-terminal helical extension (green), indicated by a green background. The presence of the C-terminal disordered protein region (red) is indicated by a yellow background, except in one nested clade, which lacks both N-terminal and C-terminal extensions, and this clade is marked by a gray background. AlphaFold predictions of proteins from each clade are used to visualize the different domains. **(B)** Phylogenetic tree of prismane domains from ACS and CODH. Blue and pink branches lead to proteins found in bacterial and archaeal genomes, respectively. Taxa with a pink background are the prismane domains found in the prismane-holding CdhCs (lower pink triangle in A). Abbreviations: acetyl-CoA synthase (ACS); CO dehydrogenase (CODH); hybrid-cluster protein (HCP).

We also identified two clades of acetyl-CoA synthases lacking both the prismane domain and the unstructured C-terminal extension. One of the clades, holding an N-terminal structured extension, branches off in between the two classic clades (CdhC and AcsB). This group of acetyl-CoA synthases is found in both bacteria (Chloroflexota, Thermodesulfobacteriota and Myxococcota) and archaea (Methanobacteria). The CODHs found in these gene clusters constitute an early branching CdhA clade. The other clade is nested inside the classical CdhC clade and represents the only clade of bacterial CdhC, although the basal branches are archaeal.

Based on our phylogenetic analysis and the previous suggestion that ACS was present in LUCA (31, 53), we propose two possible roots to the acetyl-CoA synthase phylogeny: We hypothesize that one root is in the AcsB clade and divides the bacterial AcsB from the archaeal AcsB, proposing that the ancestral ACS was fused to a prismane domain and lacked the C-terminal disordered helix. We theorize that the other root divides AcsB (ACS holding prismane domain) from CdhC-like (ACS lacking prismane domain). This root implies lateral gene transfer of AcsB between extinct bacterial lineages and Bathyarchaeota and leaves the ancestral ACS configuration ambiguous. Together, these observations reveal an unexpected structural and evolutionary plasticity of acetyl-CoA synthases across both domains of life.

## Discussion

Our structural and biochemical analyses of the ∼2 MDa ACDS supercomplex from *Methanosarcina acetivorans* contribute to the understanding of how methanogens achieve highly efficient acetyl-CoA synthesis and cleavage. This process is central to aceticlastic methanogenesis, which is estimated to produce 600-700 million tons of methane annually (15, 54), thereby contributing substantially to global greenhouse gas emissions. Rather than forming a stable bifunctional CODH/ACS complex, archaeal ACDS is built around CoFeSP and, more specifically, the N-terminal region of the CoFeSP small subunit CdhD. We show that this region promotes hexa-to octameric oligomerization and propose that the resulting CoFeSP assembly provides the scaffold for CODH and ACS association. The conservation of the CdhD and CdhA N-termini and the CdhC C-terminus across archaeal ACDS systems suggests that this scaffolded organization may be a general feature of methanogenic ACDS. In this model, CODH, ACS and CoFeSP are arranged as flexible peripheral modules connected to a central CoFeSP-based scaffold. This architecture allows catalytic modules to remain locally concentrated while retaining the conformational freedom required for acetyl-CoA synthesis and cleavage.

This organization contrasts with the bifunctional CODH/ACS complex found in bacteria, where stable CODH– ACS association is mediated by the ACS A1 domain (Fig. 6) (20, 27). In the bifunctional ACS of *Clostridium autoethanogenum*, domain A1 forms relatively stable, large interfaces with both CODH (binding interface area 1,012 Å^2^) and CoFeSP (1,625 Å^2^) (43). The absence of domain A1 in ACDS leads to a decrease of these interface areas (ACS-CODH 853 Å^2^, ACS-CoFeSP 1,218 Å^2^), suggesting that archaeal ACDS relies on weaker and more transient catalytic contacts than bacterial CODH/ACS. We propose that the ACDS oligomer compensates for this instability by keeping CODH, ACS and CoFeSP locally concentrated, while still allowing the catalytic modules to exchange binding partners.

In both ACDS and CODH/ACS, catalysis is tightly coupled to conformational dynamics of ACS (45), which in bacteria is largely enabled by the flexible linker between domains A1 and A2 (43, 55). In bacteria, ACS alternates between open and closed states: In the open conformation, the A-cluster is solvent accessible, allowing binding or release of acetyl-CoA, CoA, or CoFeSP for methyl transfer. In the closed, CODH-bound state, domain A1 and domains A2–A3 pivot to enclose the A-cluster, aligning it with and opening the CO tunnel for carbonylation or decarbonylation (27). We captured equivalent closed and open states in ACDS by cross-linking MS. As previously observed in cryo-EM structures of *Methanosarcina thermophila* CODH in complex with ACS (22), the closed state closely resembles that of bacterial CODH/ACS. The main difference is that the role of ACS domain A1 in CODH/ACS, namely CO-tunnel gating and formation of a CO alcove at the A-cluster, is taken over by CODH Rossmann-fold 2 in ACDS. The open, CoFeSP-bound state, previously observed for *Moorella thermoacetica* and *Clostridium autoethanogenum* CODH/ACS (43, 56), is described here for the first time for ACDS. In this state, protein–protein interactions are mainly established between CdhD, the N-terminal domain of CdhE and ACS domain A3, positioning the CoFeSP corrinoid domain near the A-cluster in a methyltransfer competent conformation.

**Figure 6:**
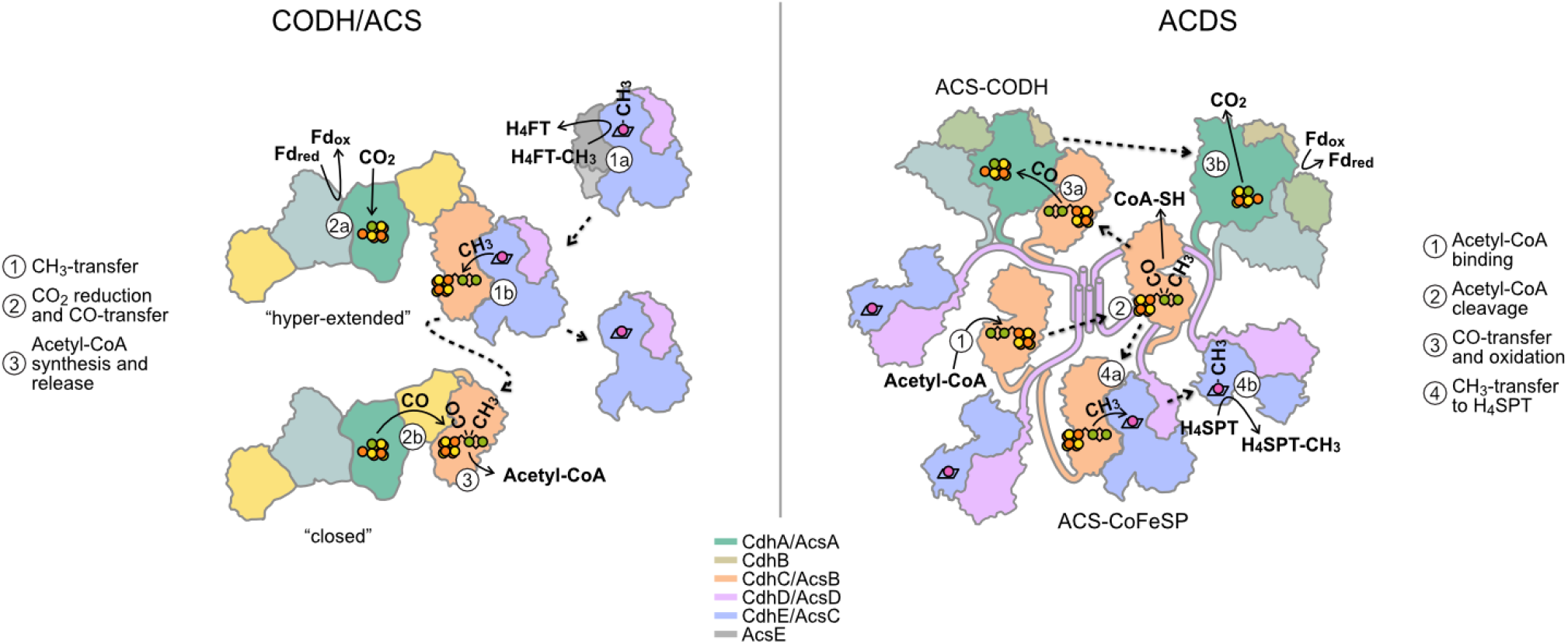
Schematic depictions of the mechanistic models of the bifunctional CODH/ACS (left) and the ACDS complex (right). CODH sub-units CdhA/AcsA and CdhB are shown in shades of green, CdhC/AcsC is depicted in orange with domain A1 colored yellow, CdhD/AcsD and CdhE/AcsC are colored pink and blue, respectively, and the methyl transferase AcsE is depicted in grey. Disordered protein regions are depicted as threads. Arrows with solid lines depict substrate/product transfers and conversions, while arrows with dashed lines depict protein conformational changes. Active site metal cofactors are shown as spheres colored by element. Abbreviations: oxidized/reduced ferredoxin (Fd_ox/red_); tetrahydrofolate (H_4_FT); tetrahydrosarcinapterin (H_4_SPT); coenzyme A (CoA); acetyl-CoA synthase (ACS); CO dehydrogenase (CODH); corrinoid iron sulphur protein (CoFeSP).

These observations raise the question of why bacteria and archaea use distinct architectures for the same chemistry. If prevention of CO leakage is not the determining factor, utilization of external CO may represent a key difference. ACS domain A1 could be advantageous for sequestering environmental CO and increasing its local concentration near the A-cluster, thereby enhancing acetyl-CoA synthesis from CO (20, 51). The two ACDS isoforms of *M. acetivorans* can catalyze either acetyl-CoA cleavage during aceticlastic methanogenesis or acetyl-CoA synthesis from CO_2_for anabolism, as well as from environmental CO during carboxidotrophic growth (34, 57, 58). In contrast, other methanogens such as *Methanosarcina barkeri* and *Methanothermobacter thermoautotrophicus* primarily use ACDS for anabolic acetyl-CoA synthesis from CO_2_or for CO oxidation during carboxidotrophic growth (59, 60). Moreover, CO is known to inhibit methanogenesis, and its primary metabolic fate in most methanogens is thought to be detoxification by CODH rather than direct utilization at the A-cluster (61), which may explain the absence of ACS domain A1 in most archaeal ACDSs. Interestingly, hybrid systems containing both CdhCtype and AcsB-type ACSs are found in some deep-sea hydrothermal vent hydrogenotrophic methanogens, like *Methanotorris formicicus* (62), *Methanothermococcus okinawensis* (63) or *Methanocaldococcus jannaschii* (64), suggesting that alternative architectures might be favored under specific environmental regimes, including high-pressure, high-temperature or low-CO conditions. For instance, it was previously speculated that a longer CO-tunnel, as extended by the A1 domain, might favor acetyl-CoA synthesis over CO oxidation under low local CO concentrations (22). The functional implications of these hybrid architectures remain to be explored.

Taken together, our findings support a model in which ACDS function is optimized through controlled structural plasticity. This organizational principle resolves the apparent paradox of a large, heterogeneous supercomplex performing tightly coordinated chemistry and provides a conceptual framework for understanding the diversity and evolution of acetyl-CoA synthase systems in anaerobic metabolism.

## Materials and Methods

### Chemicals and consumables

Unless otherwise stated, chemicals were acquired from Sigma-Aldrich/Merck, Thermo Fisher Scientific/Fisher Chemical, Carl Roth, Grüssing GmbH, VWR, Fisher Chemical, Chemsolute, Biosolute, and PanReac AppliChem (ITW Reagents).

All solvents for mass spectrometry were LC-MS-grade unless otherwise stated: water with 0.1% formic acid (Thermo, cat#85171), acetonitrile with 0.1% formic acid (ProteoChem, cat#LC6312-1L), acetonitrile (“ACN”, Fisher Scientific, cat#AA47138M6), formic acid (Thermo, cat#85178), methanol (PerkinElmer, cat#N9304938). The following reagents were used: trifluoroacetic acid (“TFA”, MS-grade, Fisher Scientific, cat#PI85183), SPE C18 disks (Empore, cat#66883-U).

### Construct design, cloning, expression, and purification of proteins

#### Cloning of CdhD fusion protein expression constructs

Residues 43 to 88 of the N-terminal disordered region (IDR) of *M. acetivorans* strain C2A ACDS Cdh2 subunit CdhD (Uniprot-ID Q8TJC2) were cloned with a C-terminal fusion to monomeric enhanced GFP (mEGFP) followed by a C-terminal Twin-Strep-II-tag, for expression in and purification from *E. coli*. The protein sequence (N’-DPMLAAALGQESAILAQHFAR-LAGMFGYPVGIGAPAAPAVSPALAA-C’) was reverse translated with *E. coli* codon usage and ordered as a gBlocks− gene fragment (IDT). The coding sequence (CDS) was amplified with primers (5’atgcaggtctcacatggatcc-tatgctggctgc3’, 5’atgcaggtctcaatcgcagctaatgccgg3’) introducing terminal overhangs with a 5’ start codon and directional BsaI recognition sites. *mEGFP* CDS was amplified from a previously cloned vector with primers (5’atgcaggtctcacgatggtgagcaagggc3’, 5’atgcaggtctcatcgacttgtacagctcg3’) introducing terminal overhangs with directional BsaI recognition sites. The inserts were cloned into backbone vector JS005 (a vector based on pET28a) using Golden Gate Assembly with BsaI-HF®v2 and T4 DNA Ligase (NEB), resulting in expression vector pEZ15. Following the same procedure, *mEGFP* CDS was amplified with primers (5’atgcaggtctcacatggtgag-caagggc3’, 5’atgcaggtctcatcgacttgtacagctcg3’) and cloned with a C-terminal Twin-Strep-II-tag into JS005, resulting in vector pEZ18.

Triple mutations of CdhD2 Leu residues 50, 57, and 64 were introduced by PCRs of pEZ15 with mutation-containing primers, followed by Gibson Assembly. Leu to Ala mutations were created with primers 64 and 65 (5’gcgctatcgccgcccaacattttgcgcgtgctgcaggcatgtttggttatc3’, 5’tgttggg-cagcgatagcgctctcttgcccagcggcagcagccagcatag3’), resulting in vector pEZ19. Leu to Asp mutations were created with primers 62 and 63 (5’gcgc-tatcgatgcccaacattttgcgcgtgatgcaggcatgtttggttatc3’, 5’tgttgggcatcga-tagcgctctcttgcccatcggcagcagccagcatag3’), resulting in vector pEZ20. Inserts of all cloned vectors were verified by Sanger sequencing (Microsynth).

#### Expression and purification of CdhD fusion constructs

CdhD fusion constructs were were expressed in *E. coli* BL21 (DE3) followed by affinity purification. Competent *E. coli* were transformed by heat-shock with vectors pEZ15, pEZ18, pEZ19, or pEZ20. Cells were cultivated in 2xYT-medium at 37 °C, 150 rpm, until reaching OD600 ∼0.8. Cultures were cooled to room temperature and expression was subsequently induced with 0.5 mM IPTG overnight at 20 °C, 150 rpm. Cells were harvested by centrifugation, flash-frozen in liquid nitrogen and stored at -80 °C until further use.

For purification, 5 g cells were resuspended in 50 mL buffer (50 mM MOPS/KOH, pH 7.2, 400 mM NaCl) containing 0.1 mM PMSF, lysozyme, DNase I (ITW Reagents), and half a tablet EDTA-free cOmplete protease inhibitor (Roche). Cells were lysed using a Microfluidizer− (Microfluidics−). Clear lysate was applied to Strep-Tactin® Sepharose (IBA Lifesciences), washed, and eluted with 2.5 mM Desthiobiotin (IBA Lifesciences). Proteins were concentrated using a 3 kDa MWCO Amicon® (Merck Millipore) and analyzed by Coomassie-stained SDS-PAGE. For Stokes radius analysis, 50 µL of purified protein were applied to a Superdex 200 Increase 3.2/300 analytical column (cytiva) mounted to an Äkta Pure FPLC in Micro kit setup (cytiva) in the same buffer. For calibration, protein from a high molecular weight calibration kit (cytiva) were used in the same buffer.

#### Expression and purification of ACDS_***Ma***_

*M. acetivorans* ACDS isoform 2 (Cdh2) was natively purified from strain MCD31, in which genes encoding Cdh1 and CdhA3 are knocked out (34). Cells were cultivated under strictly anaerobic conditions in 2 L bottles sealed with rubber stoppers, with the bottle headspace filled with 100 % N_2_. Cells were grown in modified high-salt mineral medium (65) on 120 mM sodium acetate, reduced with 0.03 g/L Na_2_S x 9 H_2_O, at 37 °C. All further steps were carried out under strictly anaerobic conditions in a vinyl anaerobic chamber (Coy Laboratory Products) filled with N_2_ and 2-5 % H_2_. Purification buffers were anaerobised by degassing, bubbling in the anaerobic chamber overnight, and addition of 2 mM DTT reducing agent. After approximately 1 week, cells were harvested in late exponential phase (OD_600_ ∼0.8) by centrifugation (13,000 x g, 30 min, RT). Cell pellets were frozen in liquid N_2_ and stored at -80 °C until further use. For cell lysis, 5 g cells were resuspended in 25 mL buffer A (50 mM potassium phosphate, pH 7.2), containing 0.2 mM PMSF, DNase I (ITW Reagents), and sonicated on ice (5x 4 min, 60 %, 0.5 s pulses). As a first purification step by HIC (hydrophobic interaction chromatography), clarified lysate was mixed with ammonium sulphate to a final concentration of 1.5 M, and loaded onto a 20 mL Phenyl Sepharose column (cytiva) using an Äkta Go FPLC (cytiva). Bound protein was washed with 0.75 M ammonium sulphate, followed by elution with a linear gradient of decreasing ammonium sulphate concentration. ACDS elution was followed by CO:Methylviologen enzymatic CODH activity, active elution fractions were pooled and diluted with buffer A to a final salt concentration of ∼200 mM. Precipitate was removed by filtration. For the second purification step by anionic IEX (anion exchange chromatography), the diluted HIC eluate was loaded onto a MonoQ 5/50 GL column (cytiva), washed with buffer A, and eluted with a linear gradient of increasing NaCl concentration. CO:MV active fractions were pooled and concentrated in a 100 kDa MWCO Amicon® (Merck Millipore). For the final purification step by SEC (size exclusion chromatography), concentrated protein was loaded onto a Superose 6 Increase 10/300 GL (cytiva) run with buffer B (50 mM potassium phosphate, pH 7.2, 400 mM NaCl). Elution fractions of the main peak were pooled and stored in the fridge in a glass vial sealed with a rubber stopper. Elution fractions were analysed by Coomassie-stained SDS-PAGE.

#### Resolution of CODH, ACS, and CoFeSP subcomplexes

ACDS subcomplexes CODH, ACS and CoFeSP were resolved under strictly anaerobic conditions by a protocol for detergent-based disassembly, previously established by Abbanat and Ferry (46), and Murakami and Ragsdale (47). Briefly, purified ACDS was diluted to ≤ 5 mg/mL, rebuffered into solubilization buffer (50 mM potassium phosphate, pH 7.2, 0.4 M NaCl, 1 % (w/v) dodecyltrimethylammonium bromide (DTAB, Tokyo Chemical Industry), 0.3 % (v/v) Triton X-100 (Merck), using a PD-10 desalting column (cytiva), and solubilized on ice for 1 h. For resolution of CODH and CoFeSP, protein was diluted with wash buffer (50 mM potassium phosphate, pH 7.2, 0.1 % (w/v) DTAB) to decrease NaCl concentration to 100 mM, which lead to precipitation of ACS. After clarification by filtration, the solubilisate was loaded onto a MonoQ 5/50 GL column (cytiva) equilibrated in wash buffer containing 100 mM NaCl. CODH and CoFeSP were separately eluted with a linear gradient of increasing NaCl concentration. To resolve ACS, protein was diluted with wash buffer to 200 mM NaCl, preventing aggregation, followed by resolution by IEX as described for the other two sub-complexes above. Detergent removal from the elution fractions was achieved by SEC using a Superose 6 Increase 10/300 GL (cytiva) for ACS, or by buffer exchange using a PD-10 desalting column (cytiva) for CODH and CoFeSP into buffer B (50 mM potassium phosphate, pH 7.2, 400 mM NaCl. Proteins were concentrated using 30 kDa MWCO Amicons® (Merck Millipore) and analyzed by Coomassie-stained SDS-PAGE.

### Acetyl-CoA formation and CO-competition assays

Kinetics of acetyl-CoA formation by and CO leakage from the purified ACDS complex were assessed by colorimetric assays as previously described (51). Purified ACDS was buffer-exchanged into assay buffer (100 mM MOPS/NaOH, pH 7.2, 400 mM NaCl) to remove DTT using Zeba− Spin Desalting Columns (Thermo Scientific). Assays were measured in 1 mL solution in Quartz cuvettes sealed with rubber stoppers to maintain a closed headspace of 0.4 mL; kinetics were recorded over 20 min at 37 °C using a NanoPhotometer® NP80 (implen) recording a full spectrum from 200-900 nm every 30 s. Assays were incubated at 37 °C for several minutes before adding ACDS right before the start of the measurement. All assay mixtures contained the following components diluted from stock solutions in assay buffer: 5 mM NaHCO_3_, 0.3 mg/L bovine carbonic anhydrase (Merck), 5 mM Ti(III)-EDTA; Ti(III)-EDTA was prepared as previously described (66) from 30 % Ti(III)Cl_3_ (Sigma-Aldrich) and stored anaerobically, protected from light and in the fridge prior to the start of the experiment. To assess acetyl-CoA formation, 200 µM CoA (MP Biomedicals), 50 µM methyl-Co(III)binamide (synthesis described in the supplementary information) and 0.5 µM (relative to the protomer) ACDS were added. To assess CO leakage, 7 µM (relative to the protomer) bovine hemoglobin (Merck) was added in addition. Negative controls contained all components except ACDS. Concentrations of products Co(I)binamide and carboxyhemoglobin were calculated from absorbances at 465 nm and at 420 and 431 nm, respectively, using a method and molar extinction coefficients previously published (66, 67). Assays were carried out as technical duplicates or triplicates and as biological duplicates.

### Mass photometry

Measurements were conducted using a TwoMP mass photometer (Refeyn Ltd., UK), switched on one hour before the first measurement. Microscope slides (1.5 H, 24 × 60 mm, Carl Roth, Germany) and silicon gaskets (CultureWellTM, CW-50R-1.0, 50-3 mm diameter x 1 mm depth) were prepared by thorough rinsing with isopropanol, followed by rinsing with Milli-Q water, and drying in a stream of pressurised air. Immersion oil (IMMOIL-F30CC, Olympus, Japan) was used. For data collection, the software AcquireMP (Refeyn Ltd., v. 2023 R1.1) was used. Sample was kept anaerobic right before starting the measurement by dilution in buffer (50 mM potassium phosphate/MOPS, pH 7.2, 400 mM NaCl) to 12.5-50 nM. Data processing and analysis was carried out with the software DiscoverMP (Refeyn Ltd., v. 2023 R1.2). For calibration, a homemade protein standard was used (86-430 kDa).

### Microfluidic modulation spectroscopy

Infrared spectra of CdhD fusion construct CdhD_IDR_-mEGFP-TS (pEZ15) and mEGFP-TS (pEZ18) were recorded with an Aurora (RedShiftBio) using protein concentrations of 0.9 and 1.1 mg/mL, respectively. To ensure buffer matching, proteins were freshly buffer exchanged into 50 mM MOPS, pH 7.2, 400 mM NaCl by gel filtration using a Superdex 200 Increase 10/300 GL (cytiva). Samples were measured in a 96-well round-bottom plate (Corning) sealed with Zone-Free− sealing film (Excel Scientific). Each sample was measured in triplicate, and normalized average absolute absorbance spectra as well as second derivative spectra were generated using the delta software (RedShiftBio). For the absolute absorbance spectra, a nominal fit displacement factor of 0.6 was applied over a fitting range of 1720-1680 cm^−1^. Second derivative spectra were processed using Savitzky-Golay smoothing with a window size of 19 wavenumbers.

### Shotgun proteomics

A 50 µg aliquot of the purified protein was reduced at 90 °C for 10 minutes using DOC (deoxycholate) in ammonium bicarbonate (pH 8.2) and TCEP (tris(2-carboxyethyl)phosphine). It was then incubated for 30 minutes at 25 °C with iodoacetic acid (IAA) and subsequently digested overnight at 30 °C with MS-approved trypsin (Serva). Before LC–MS analysis, samples were desalted using C18 microspin columns (Nest Group) according to the manufacturer’s instructions. The dried and reconstituted peptides were then analysed by liquid chromatography–mass spectrometry, performed on an Orbitrap Exploris 480 instrument connected to an Ultimate 3000 RSLC nano and a Nanospray ion source (all from Thermo Scientific). Peptide separation was carried out on a reverse-phase HPLC column (75 μm × 40 cm) internally packed with C18 resin (2.4 μm; Dr. Maisch) using a 30-minute gradient (formic acid/acetonitrile). The MS data were aligned against an in-house *Methanosarcina acetivorans* protein database using Sequest, which is integrated into the Proteom Discoverer 1.4 software (Thermo Scientific). MS data were searched against an in-house *Methanosarcina acetivorans* protein database using Sequest embedded into Proteom Discoverer 1.4 software (Thermo Scientific).

### Inductively coupled plasma mass spectrometry analysis

ICP-MS was used to measure metal concentrations in biological duplicates of natively purified ACDS. Briefly, acid digestion of 315 µg sample (12.6 mg/mL protein, determined by BCA assay) or of 540 µg sample (3 mg/mL, determined by Bradford) was performed in 11% (v/v) HNO_3_ (Suprapur, Merck), for 3 h at 80 °C. This was subsequently diluted 5.5 x with Chelex-treated water to a total volume of 250 µL per sample. Internal standards, indium (2 ppb) and germanium (20 ppb) (both from VWR Merck), were added to each sample for normalization. Elemental analysis was conducted using an inductively coupled plasma mass spectrometer iCAP Q (ICP-MS, Thermo Fisher Scientific), equipped with an SC4DX autosampler (Elemental Scientific) and a MicroFlow PFA-100 nebuliser. Quantification of metals was done by comparison with serial dilutions of the ICP multi-element standard solution XVI (Merck). The ICP-MS operated with a reaction cell using a helium/hydrogen gas mixture (93/7%). Data acquisition was performed in triplicate using Qtegra software v2.18 (Thermo Fisher Scientific). Blank values and quality thresholds were calculated using protein-buffer standards. The measured concentrations (initially in ppb) were converted to molar units (µM) of metal per sample for quantitative analysis.

### Crosslinking mass spectrometry

#### Protein crosslinking

Proteins were crosslinked under strictly anaerobic conditions in an anaerobic chamber, as described above. Crosslinking buffer (50 mM potassium phosphate, pH 7.2, 400 mM NaCl) and MiliQ water were anaerobised by degassing and bubbling in the anaerobic chamber overnight, without addtion of DTT. Purified ACDS was buffer exchanged into crosslinking buffer using SEC (Superse 6 Increase 10/300 GL, cytiva) or PD columns (cytiva), concentrated to 4.6-10 mg/mL. Sulfo-SDA (Thermo Scientific Pierce) was freshly dissolved to 20 mM (7.06 mg/mL) in anaerobic MiliQ water and immediately added to the protein at 1:0.38, 1:0.5, 1:075, 1:1 and 1:2 protein to crosslinker mass ratio. Reactions were incubated light-protected at room temperature for 30 min. Next, the mixtures were transferred into 1.5 mL tube lids and irradiated with a UV handheld lamp at 365 nm (ATG Kriminaltechnik) for 30 min on ice. Reactions were quenched with 100 mM Tris/HCl, pH 7.2, denatured and separated by Coomassie-stained SDS-PAGE. Protein bands corresponding to crosslinked, non-aggregated protein, were excised and washed intensively with 70 % (v/v) 50 mM NH_4_HCO_3_. 30 % (v/v) acetonitrile to remove Coomassie, before gel pieces were dried with 100 % acetonitrile.

#### Peptide size-exclusion chromatography

Dried peptides were eluted from StageTips before being resuspended in 25 µL of 30% acetonitrile with 0.1% TFA, vortexed, and sonicated for 1 min before injecting onto a Superdex 30 Increase 10/300 GL column (Cytiva, cat#GE29219757) with a 20 µL injection loop using an Äkta pure micro system and eluting peptides using the following gradient: 0–4.5mL (150μL/min), 4.5–9.5mL (250μL/min), 9.5–14.5mL (300μL/min), 14.5– 28.5mL (400μL/min). Crosslinked peptides eluted over the range 9.5– 14.5mL as 250 μL fractions into protein lo-bind tubes. Samples were dried *in vacuo* and subsequently desalted via C18 StageTips before peptides were eluted into lo-bind tubes to dry.

#### Crosslinking mass spectrometry

Dried peptides were resuspended in 10 μl of 1.6% acetonitrile with 0.1% formic acid, vortexed, and sonicated for 1 min before injecting 1 μg of estimated peptide sample onto a Thermo Eclipse Orbitrap coupled to a Vanquish Neo HPLC system. Peptides were ionized using an EASY-Spray source and eluted over an EASY-Spray PepMap Neo 75 μm × 500 mm C18 column with LC–MS quality water or acetonitrile with 0.1% formic acid (mobile A and B, respectively). The flow rate was 0.3 μl per minute using gradients optimized for each chromatographic fraction from offline fractionation, ranging from 1.6% mobile phase B to 44% mobile phase B over 95 min, followed by a linear increase from 44% to 76% mobile phase B in 2.5 min, respectively.

Peptides were analyzed using the following MS global parameters: method duration of 95 min; infusion mode, liquid chromatography; expected LC peak widths, 30 s; advanced peak determination checked; default charge state of 2; EASY-IC internal mass calibration; NSI ion source; static spray voltage at 2,000 V in positive mode; static gas mode with a sweep gas setting of 2; ITT temperature of 280 °C. Samples were collected using the following shared scan parameters: duty cycle of 3 s; MS-OT at 240,000 resolution; normal mass range; quadrupole isolation checked; scan range of 380–2,000; RF lens of 35%; custom AGC target of 150% with a maximum injection time of 100 ms; one microscan in profile mode at positive polarity; EASY-IC checked; subbranch MIPS, peptide; subbranch intensity, 2.5 × 10^4^; subbranch charge state 3–7; subbranch dynamic exclusion of one time after 30 s with a mass tolerance of 10 ppm; exclude isotopes and within cycle checked and dependent scan on single charge state unchecked; subbranch priority 1: subbranch A: charge state, 4; precursor selection range, 380–1,800; subbranch B: charge state, 5; precursor selection range, 380–1,350; subbranch C: charge state, 6–7; precursor selection range, 380–1,000; rejoined subbranch: sort by intensity – most intense; subbranch ddMS2 OT HCD: isolation mode, quadrupole with a window of 1.4 *m*/*z*; stepped HCD (normalized) of 20, 26 and 30; detector type, Orbitrap at 60,000 resolution from 150–2,000 *m*/*z*; AGC target of 750% with a maximum injection time of 250 ms; one microscan; centroid data. Subbranch priority 2: subbranch: charge state, 3; precursor selection range, 380–2,000; subbranch: sort by intensity – most intense; subbranch ddMS2 OT HCD: isolation mode, quadrupole with a window of 1.4 *m*/*z*; stepped HCD (normalized) of 20, 26 and 30; detector type, Orbitrap at 60,000 resolution from 150–2,000 *m*/*z*; AGC target of 750% with a maximum injection time of 250 ms; one microscan; centroid data.

If enough material was available, a secondary “BoxCar” acquisition was performed using the global MS parameters with the following changes: duty cycle of 5.12 s, OT tSIM with 10 max multiplexed ions (user-defined, see table below), SIM Window Mode – Center Mass, auto maximum injection time, unscheduled time mode; subbranch A: charge state, 2–6. The following windows were utilized for both BoxCar tSIMs:

| Compound | Center Mass (m/z) | z | MSX ID | Isolation Window (m/z) | Normalized AGC Target (%) |
| --- | --- | --- | --- | --- | --- |
| Boxcar1-1 | 414.1 | 1 | 1 | 30.2 | 50 |
| Boxcar1-2 | 462.2 | 1 | 1 | 24.8 | 50 |
| Boxcar1-3 | 504.1 | 1 | 1 | 23.6 | 50 |
| Boxcar1-4 | 545.35 | 1 | 1 | 23.1 | 50 |
| Boxcar1-5 | 587.8 | 1 | 1 | 25 | 50 |
| Boxcar1-6 | 634.05 | 1 | 1 | 26.7 | 50 |
| Boxcar1-7 | 686 | 1 | 1 | 31.4 | 50 |
| Boxcar1-8 | 748.45 | 1 | 1 | 38.9 | 50 |
| Boxcar1-9 | 829.65 | 1 | 1 | 48.5 | 50 |
| Boxcar1-10 | 957.2 | 1 | 1 | 91.6 | 50 |
| Boxcar2-1 | 439.5 | 1 | 2 | 26.6 | 50 |
| Boxcar2-2 | 483.45 | 1 | 2 | 23.7 | 50 |
| Boxcar2-3 | 524.85 | 1 | 2 | 23.9 | 50 |
| Boxcar2-4 | 566.15 | 1 | 2 | 24.3 | 50 |
| Boxcar2-5 | 610.5 | 1 | 2 | 26.4 | 50 |
| Boxcar2-6 | 658.85 | 1 | 2 | 28.9 | 50 |
| Boxcar2-7 | 715.35 | 1 | 2 | 33.3 | 50 |
| Boxcar2-8 | 786.65 | 1 | 2 | 43.5 | 50 |
| Boxcar2-9 | 882.65 | 1 | 2 | 63.5 | 50 |
| Boxcar2-10 | 1101 | 1 | 2 | 202 | 50 |

If enough material was still available, additional, windowed acquisitions were performed using the global MS parameters with the following changes: dynamic exclusion of one time after 7.5 s with a mass tolerance of 10 ppm; precursor selection range, 380–605 or 600–805 or 800–1,005 or 1,000–1,600 (different for each injection).

A recalibration of the precursor *m*/*z* was conducted on the basis of high-confidence (<1% FDR) linear peptide identifications for the following protein sequences of CdhA2, CdhB2, CdhC2, CdhD2, and CdhE (Uniprot-IDs Q8TJC6, Q8TJC5, Q8TJC4, Q8TJC2, and Q8TH44, respectively). To identify crosslinked peptides, the recalibrated peak lists were searched against the sequences and the reversed sequences (as decoys) of crosslinked peptides using the Xi software suite (v.1.8.6) (68). The following parameters were applied for the search: MS1 accuracy = 2 ppm; MS2 accuracy = 5 ppm; enzyme = trypsin, allowing up to three missed cleavages and two missing monoisotopic peaks; crosslinker A = SDA, with an assumed NHS-ester reaction specificity for lysines, serines, threonines, tyrosines, and protein N-termini crosslinked to the diazirine (A/C/D/E/G/H/I/K/L/P/S/T/V/Y/protein C-termini /protein N-termini); crosslinker B = Ccm-SDA, with an assumed NHS-ester reaction specificity for lysines, serines, threonines, tyrosines, and protein N-termini crosslinked to cysteine (adjusted for the fixed modification mass for cysteines containing the carbamidomethyl modification) for the diazirine; fixed modifications = carbamidomethylation on cysteine; variable modifications = oxidation on methionine, pyroglutamate (N-terminal to Q, -17.026549105), hydrolyzed SDA (K/S/T/Y/protein N-termini, 100.05243), loop-linked SDA (K/S/T/Y/protein N-termini, 82.04186484; linear modifications = deamidation (N, 0.984016), unreacted SDA (K/S/T/Y/protein N-termini, 110.04801284). MS cleavage of SDA crosslinks was considered during searches. The CSM candidates were then filtered to a 2% FDR at the residue-pair level using xiFDR (v.2.2.1) (69), and visualized and analyzed in xiVIEW (70).

### AlphaLink2 modelling

CODH-ACS, CoFeSP-ACS, CoFeSP-CODH, and CODH-ACS-CoFeSP protein-protein interactions based on crosslinks were predicted using AlphaLink2 (71). As inputs, all identified heteromeric crosslinks and one copy per chain of full-length protein sequences of CdhA2, CdhB2, CdhC2, CdhD2, and CdhE (Uniprot-IDs Q8TJC6, Q8TJC5, Q8TJC4, Q8TJC2, and Q8TH44, respectively) were used. 11 crosslinks were used for CODH-ACS prediction, 22 crosslinks were used for CoFeSP-ACS prediction, 5 crosslinks were used for CoFeSP-CODH prediction, and 55 crosslinks were used for CODH-ACS-CoFeSP prediction. Ten models were predicted per interaction and ranked by pTM scores. Figures were prepared using open-source PyMOL (Schrodinger, LLC. 2010. The PyMOL Molecular Graphics System, Version 3.0.0).

### Small-angle x-ray scattering

BioSAXS experiments were conducted at beamline BM29 at the ESRF (European synchrotron radiation facility) equipped with a Pilatus3 2M in vacuum detector (72). All samples were measured at a wavelength of 1 Å and a q-range of 0.025-6 nm^−1^. For batch SAXS measurements, ACDS protein was purified and stored anaerobically in anoxic buffer (50 mM potassium phosphate, pH 7.2, 400 mM NaCl, 2 mM DTT), buffer matching was ensured by SEC. A dilution series from 10 mg/mL to 0.5 mg/mL in the same buffer was prepared aerobically right before the experiment. Samples were automatically injected into and measured in a 1 mm diameter quartz capillary. Before and after each sample, the buffer was measured. Ten 1 s frames were recorded, frames selected for processing were averaged and averaged buffer frames were subtracted. For the SEC-SAXS measurement, purified ACDS protein was buffer-exchanged into another buffer (50 mM MOPS, pH 7.2, 400 mM NaCl, 2 % (w/v) sucrose, 2 mM reduced glutathione, 2 mM DTT (both added after buffer anaerobisation), 2 mM NaNO_3_, 2 mM ascorbic acid) to decrease radiation damage, using SEC (Superose 6 Increase 10/300) to ensure buffer matching. 100 µL sample at 5 mg/mL protein concentration were loaded onto a Bio SEC-3, 300 Å column (Agilent) connected to a Shimadzu HPLC; eluate was measured in a 1 mm diameter quartz capillary; a flow-rate of 0.2 mL/min was used, 500 frames of 3 s were recorded. Data were processed with CHROMIXS (73) and PRIMUS from the ATSAS software package (v. 4.1.1) (74) and ScÅtter IV (75).

### Cryo-EM single-particle analysis

#### Grid preparation

For cryo-EM SPA, QUANTIFOIL R 1.2/1.3 copper grids were glow discharged for 25 s with a current of 15 mA in a PELCO easiGlow device (Ted Pella). Natively, anaerobically purified ACDS protein complex (1 mg/mL in 50 mM KH_2_PO_4_/K_2_HPO_4_, pH 7.2, 150 mM NaCl) was freshly crosslinked with 1 mM BS^3^ (Thermo Scientific) for 15 min, quenched by adding 50 mM Tris/HCl, pH 7.0, supplemented with 0.04 % (w/v) n-Octyl-beta-D-gluco-pyranoside, and vitrified in an anaerobic chamber (Coy Laboratory Products) filled with 95 % N2, > 1.5 % H2, using a Vitrobot Mark IV (Thermo Fisher Scientific). 4 µL protein were applied onto the grid, blotted with blot force 4 for 4 s at 4 °C and 100 % humidity, and automatically plunged into 37 % (v/v) liquid ethane (in propane).

#### Data collection and processing

Data was acquired using SerialEM software on a Jeol CryoArm200 microscope operated at 200 kV, equipped with a K2 Summit direct electron detector, with x60,000 magnification, 0.85 Å pixel size, with a total dose of 50 e-/Å2. 5,987 micrographs were acquired.

In CryoSPARC (76), the micrographs were motion corrected using patch motion correction (77), and contrast transfer functions (CTFs) were estimated using patch CTF estimation. For processing of ACDS particles, particles were initially picked with blob picker, followed by manual particle curation and two rounds of 2D classification and selection, resulting in 99,209 particles extracted with a box size of 550 px. One ab initio and four decoys were reconstructed and used for heterogeneous refinement, resulting in five maps of which four were further pursued and refined with homogeneous refinement. Class 3 with 20,624 particles and a nominal resolution of 19.59 Å (gsFSC) was further pursued. Homogeneous refinement applying different symmetries (C2, C3 C4, D2, D3, D4, T), local refinements and 3D classifications with masks around outer shell blobs, 3D variability analysis (78) and 3D flex analysis (79) did not result in improved resolution. Particles were downsampled by Fourier cropping to 190 px and converted to Relion star files using cs for processing in Relion 5.0 (80). Class 3 was further refined using auto-refine and 10 3D classes were obtained. 3D multi-body refinement (81) with 14 masks engulfing the 14 main blobs was attempted, but resulted in local maps with increased noise.

For processing of CODH particles, particles were initially picked with blob picker, manually curated and subjected to 2D classification and selection, resulting in 110,853 particles. The particles were used for two rounds of Topaz train, extract, 2D classification and selection, resulting in 147,351 particles. These particles were re-extracted with a box size of 360 px using only micrographs with CTF fit resolution ≤ 5 Å, resulting in 77,949 particles. From these particles, one ab-initio map was reconstructed and refined with C2 symmetry using homogeneous refinement. The map was used to create 50 2D templates for template picking on the previously curated set of micrographs, which resulted in 371,631 particles after manual inspection of particle picks. Particles were extracted with a box size of 360 px, three ab-initio maps were reconstructed and refined with heterogeneous refinement. The map representing CODH was refined with C2 symmetry and a mask around CODH using homogeneous refinement, non-uniform refinement (82) with optimizing per-particle defocus, fitting tilt and trefoil, and with local refinement, resulting in a map with 3.55 Å resolution (0.143 GSFSC). The map was sharpened with a B-factor of 116.2. For CODH location analysis, particle locations of the 147,351 CODH particles from curated blob picking, and 124,730 ACDS particles from four best 3D reconstructions after heterogeneous refinement were exported to Relion star files. Particle locations were used to calculate and plot the distances between centres of CODH and ACDS particles within 1,000 px maximum distance. The CODH particle population within 180 px (153 Å) distance from ACDS particles was selected, resulting in 32,239 particles. These particles were imported into Relion 5.0 (80) and used for 3D ab-initio reconstruction followed by 3D classification with 5 classes. Class 1 (4,356 particles) and class 3+4 (14,637 particles) were further refined with masks around their CODH, or CODH with attached blob.

#### CODH model refinement

As initial model, an AlphaFold2 (38) prediction of *M. acetivorans* CODH (CdhA_2_B_2_) was used. The model was first fitted into the map using ChimeraX (83) and non-resolved protein regions were truncated in WinCoot (v. 0.9.8.93 and 0.9.8.96) (84). WinCoot and PHENIX (v. 1.19.2-4158 and 2.0-5936) real-space refinement (85) were iteratively used to refine the model, with secondary structure restraints, NCS constraints, and using reference model restraints from *Methanosarcina thermophila* CODH (pdb 9c0r) (22). Metal clusters were first manually fitted and included in refinement. Molecular graphics and analyses were performed with UCSF ChimeraX (83), developed by the Resource for Biocomputing, Visualization, and Informatics at the University of California, San Francisco, with support from National Institutes of Health R01-GM129325 and the Office of Cyber Infrastructure and Computational Biology, National Institute of Allergy and Infectious Diseases.

### ACDS integrative modelling

To visualize conformational states of the expected ACDS structural ensemble in 6- and 8-fold stoichiometry, the integrative modelling software Assembline (50) was used. The consensus cryo-EM map was gaussian filtered with 10 standard deviations and Fourier cropped to a pixel size of 11.7 Å, to remove artificial high-resolution features. Input models for *M. acetivorans* CODH (CdhA_2_B_2_), ACS (CdhC) in open and closed conformation, and CoFeSP (CdhDE) were created from the predicted AlphaLink2 models, and only high-confidence regions without the terminal disordered regions were used as rigid bodies (residues CdhA 41-805, CdhB 2-170, CdhC 2-400, CdhD 91-436, CdhE 2-468). Furthermore, an AlphaFold2 model of 8-fold CdhD coiled coil (CdhD residues 44-70) was used. Fit libraries of each rigid body within the cryo-EM map were calculated using Efitter, with 100,000 searches, accepting fits with 60 % of the input model within the map envelope. The fit libraries were used for simultaneous fitting of multiple rigid bodies with simulated annealing Monte Carlo simulation, running 5,000 simulations with global optimization; 4 CODH, 8 CoFeSP, 4 ACS in open and 4 ACS in closed conformation, and 1 8-fold CdhD coiled coil were used, 90 hetero- and homomeric overlapping crosslinks in between CdhA-CdhC, CdhD-CdhC, CdhA-CdhD, and CdhA-CdhE were used, CoFeSP rigid bodies distance to coiled coil were used as connectivity restraints, the CdhD missing residues were modelled as 10-residue resolution spheres, and excluded volume restraint was included in the scoring function to minimize clashes. Next, a narrowed down fit library of rigid bodies was created from the top 5 % scoring models, and used for recombination, running 35,000 simulations with the same restraints used for global optimization. The top models after recombination contained obvious subunit clashes. Rigid bodies were manually refined in the top scoring model to remove major clashes, and the resulting rigid bodies were used as input for refinement, running 3 simulations. For refinement, an extended set of 78 heteromeric and 23 homomeric overlapping crosslinks was used; Flexible residues within regions found in crosslinks (CdhD 71-90, CdhA 15-40, CdhC 401-465) were modelled as single-residue beads and used as connectivity restraints; The weight of the excluded volume restraint was increased to prevent the formation of clashes, and refinement was run excessively (16.3 * 10^6^ simulated annealing steps) to optimize restraints. The top scoring model was reconstructed including the modelled flexible residues, cofactors were added by superposition with respective structures (pdb-id 9c0s for A- and C-clusters, 2ycl for CdhE corrinoid and [4Fe-4S]), and geometric issues in the resulting model were resolved using ISOLDE (86) and phenix.dynamics (85). A SAXS profile of the model was simulated using FoXS (87) and the model was analyzed and visualized using xiVIEW (70), ATSAS (74), and open-source PyMOL (Schrodinger, LLC. 2010. The PyMOL Molecular Graphics System, Version 3.0.0).

### Phylogenetics

Amino acid sequences of ACS, CODH and HCP were sampled from the joint genome institute’s (JGI) integrated microbial genome & metagenomes database (IMG) (88) and NCBI RefSeq (89) using BLAST (90) Predicted structures of the proteins were either found in the AlphaFold Database (91) or folded using the AlphaFold3 Server (92). The predicted structures were translated into 3Di characters and aligned using Foldmason/Foldseek (93) and mafft-linsi (94) using the 3Di scoring matrix from Foldseek. Maximum-likelihood phylogenies were constructed using iqtree (95, 96). For the partitioned ACS tree, the 3Di part was inferred under the substitution matrix substitution model described here (97), with invariable sites and FreeRate model, whereas the Le-Gascuel (LG) substitution matrix substitution model and FreeRate model was used for the amino acid part, as determined by automatic best-fit evolutionary model selection. The phylogeny of prismane domains was inferred using 3Di only, as described for ACS above. The robustness of the phylogenies was assessed using 1000 ultrafast bootstraps.

## Data availability

The cryo-EM maps were deposited to the EM Data Bank under accession codes EMD-58600 (ACDS complex) and EMD-58601 (CODH subcomplex). The atomic models will be deposited to the PDB-IHM (integrative model of the ACDS complex) for which an accession code will be provided, and to the PDB under accession code 31OX (cryo-EM structure of the CODH sub-complex). The mass spectrometry proteomics data have been deposited to the ProteomeXchange Consortium via the PRIDE (98) partner repository with the dataset identifier PXD078296 and 10.6019/PXD078296. The SEC-SAXS data has been deposited to the SASBDB (99) under accession code SASD2C2.

## Acknowledgements

We thank Michael Rother from Dresden University of Technology for providing us with the *M. acetivorans* strain essential for this study. We acknowledge the contributions of the cryo-EM Facility of the Philipps-University Marburg and are grateful to Anuj Kumar to assist in cryo-EM data collection. We would like to thank the staff of the ESRF and EMBL Grenoble, in particular Mark Tully and Stephanie Hutin, for assistance and support in using beamline 29 under proposal numbers MX-2584 and MX-2685. We thank Jan Schäfer from RedShiftBio for acquiring and processing MMS data. We thank Marburg University for financially supporting the ChemBio research-based platform. This research was supported by the Intramural Research Program of the National Institutes of Health, National Cancer Institute (NCI), Center for Cancer Research, Project number ZIA BC012114-03 (F. J. O’Reilly). This project has been funded in part with Federal funds from the Frederick National Laboratory for Cancer Research, National Institutes of Health, under contract HHSN261200800001E (F. J. O’Reilly). The contributions of the NIH authors were made as part of their official duties as NIH federal employees, are in compliance with agency policy requirements, and are considered Works of the United States Government. However, the findings and conclusions presented in this paper are those of the authors and do not necessarily reflect the views of the NIH or the U.S. Department of Health and Human Services. We acknowledge and thank the American Lebanese Syrian Associated Charities (ALSAC) for financial support (F. J. O’Reilly). This work was supported by the European Union’s Horizon 2020 research and innovation programme (Two-CO2-One; grant agreement no. 101075992 to J.M.S.) and by the Deutsche Forschungsgemeinschaft (DFG, German Research Foundation) – RTG 2937 (project number 505997786) to J.M.S. The views and opinions expressed are those of the author(s) only and do not necessarily reflect those of the European Union or the European Research Council. Neither the European Union nor the granting authority can be held responsible for them. This paper was typeset with the bioRxiv word template by @Chrelli: www.github.com/chrelli/bioRxiv-word-template

## Author contributions

E.J.Z. and T.R.-T. designed experiments, purified proteins and processed cryo-EM data. E.J.Z. cloned expression constructs, prepared samples for crosslinking MS, conducted biophysical experiments and analyzed data (SAXS, MP, SEC) and built structural models. T.R.-T. prepared cryo-EM grids. A.C. collected and processed crosslinking MS data. S.A. conducted the phylogenetic analysis. J.K. did shotgun proteomics for protein identification. D.D. conducted ICP-MS experiments. F.A. and O.V. synthesized methylcobinamide. G.K.A.H., F.J.O’R. and J.M.S. conceptualized and supervised the research. E.J.Z. and J.M.S. wrote the original draft of the paper. All authors participated in the discussion and paper editing. J.M.S. and T.R.-T. originally conceptualized the project.

