## Extended Data for "Interface swapping orchestrates carbon transfer in the archaeal acetyl-CoA decarbonylase/synthase"

bioRxiv

[1] Philipps-University Marburg, Department of Chemistry and Center for Synthetic Microbiology (SYNMIKRO), Karl-von-Frisch-Strasse 14, Marburg, Germany
[2] Center for Structural Biology, Center for Cancer Research, National Cancer Institute, National Institutes of Health, Frederick, MD, USA
[3] Department of Biology, Philipps-University Marburg, Marburg, Germany
[4] Max Planck Institute for Terrestrial Microbiology, Marburg, Germany
[5] Department of Environmental Biotechnology, Helmholtz Centre for Environmental Research (UFZ), Leipzig, Germany
[6] Philipps-University Marburg, Department of Chemistry, Marburg, Germany
[7] Center of Excellence for Data Driven Discovery, Department of Structural Biology, St. Jude Children's Research Hospital, Memphis, Tennessee 38105, United States
[8] Microbes-for-Climate (M4C) Cluster of Excellence, Marburg, Germany
[†] These authors contributed equally

### The PDF file includes:

Extended Data Figures 1-9

Extended Data Table 1

### Extended Data Figures

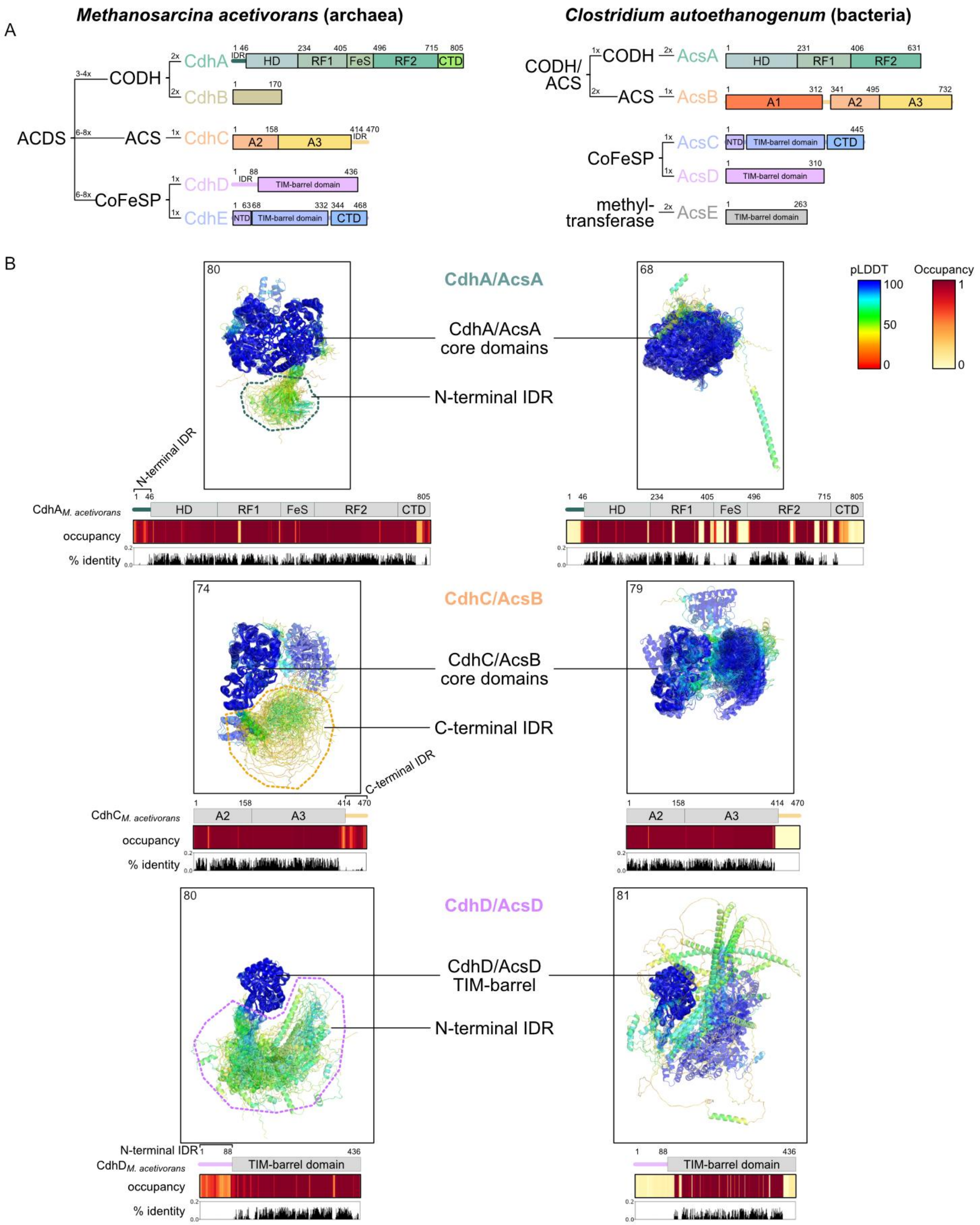

**Extended Data Fig. 1: Subunits of the bacterial and archaeal CODH/ACS/ACDS enzyme complexes. (A)** Schematic representation of the five archaeal and bacterial subunits, CdhA-E and AcsA-E, respectively, and the subcomplexes they form; domains are depicted as rectangles, unstructured regions as threads. **(B)** Phylogenetic analysis of CdhA/AcsA, CdhC/AcsB, and CdhD/AcsD terminal disordered regions. The analyzed dataset of protein sequences was obtained from a previous publication (1). AlphaFold2 models were aligned to the M. acetivorans cdh2 homologs, as references, with foldmason (2), coloured by pLDDT score, and are depicted as transparent cartoon; the No. of sequences/models compared is noted in the top left corner of each box. Structurally aligned amino acid sequences were analyzed with Jalview (3) and occupancy and % identity plots created with Biopython (4). As reference, schematics of the M. acetivorans cdh2 homologs' domain architectures are shown above the plots. Abbreviations: acetyl-CoA decarbonylase/synthase (ACDS); acetyl-CoA synthase (ACS); CO dehydrogenase (CODH); corrinoid iron sulfur protein (CoFeSP); N-terminal helical domain (HD); Rossmann-fold domain (RF); iron-sulfur domain (FeS); C-terminal domain (CTD); N-terminal domain (NTD); intrinsically disordered protein region (IDR).

A

| Accession | Description | Score | Coverage | # Unique Peptides | # Peptides | # PSMs |
|---|---|---|---|---|---|---|
| Q8TJC6 | CdhA2 | 3443.2 | 87.83% | 14 | 90 | 1670 |
| Q8TJC2 | CdhD2 | 2618.01 | 84.86% | 28 | 38 | 966 |
| Q8TJC4 | CdhC2 | 2067.69 | 79.36% | 23 | 58 | 1131 |
| Q8TH44 | CdhE | 1537.77 | 76.50% | 39 | 55 | 892 |
| Q8TJC5 | CdhB2 | 477.73 | 94.71% | 18 | 21 | 274 |

B

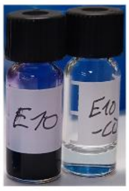

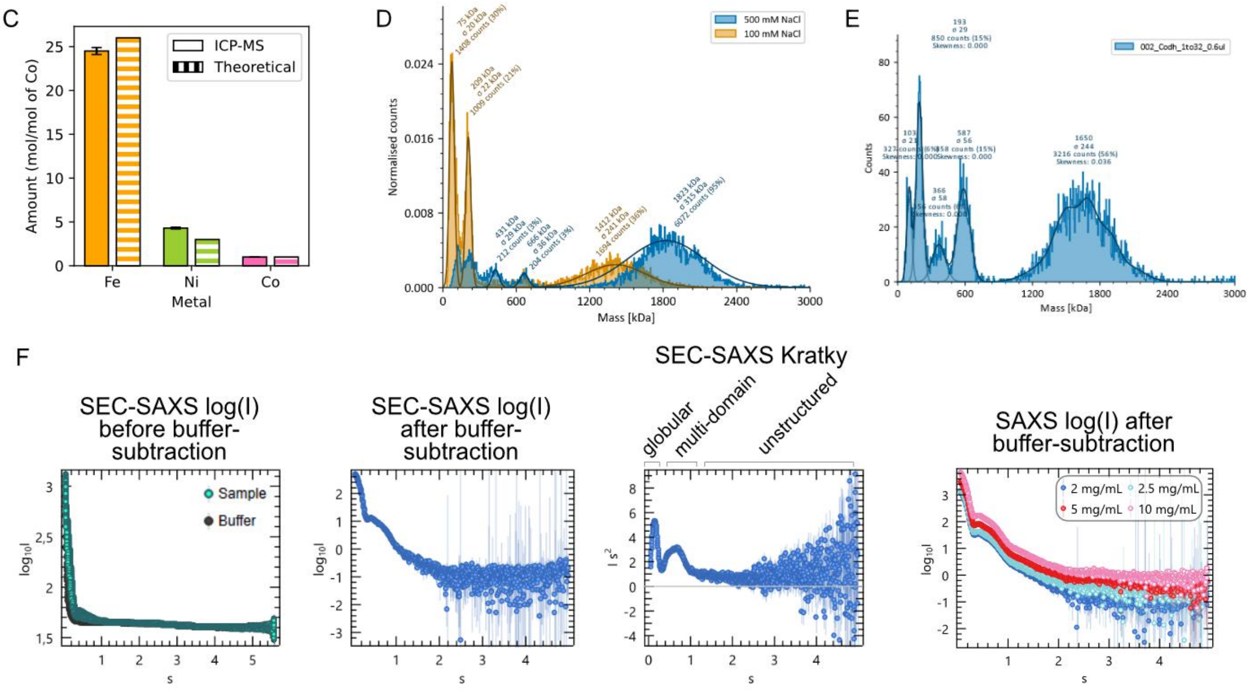

**Extended Data Fig. 2: Native purification of M. acetivorans ACDS Cdh2. (A)** Protein LC-MS results of the five ACDS subunits in the purified protein sample. **(B)** Qualitative CO:Methylviologen colorimetric assay of ACDS IEX fraction with and without the addition of CO in the headspace. **(C)** ICP-MS results of iron (Fe), nickel (Ni) and cobalt (Co), normalised to 1 Co; the theoretically expected quantities were calculated assuming equal stoichiometry of the five subunits with full metal cofactor occupancy. **(D)** Mass photometry in buffer with high (500 mM) and low (100 mM) salt (NaCl). **(E)** Mass photometry carried out in an anaerobic chamber. **(F)** SEC-SAXS and SAXS results.

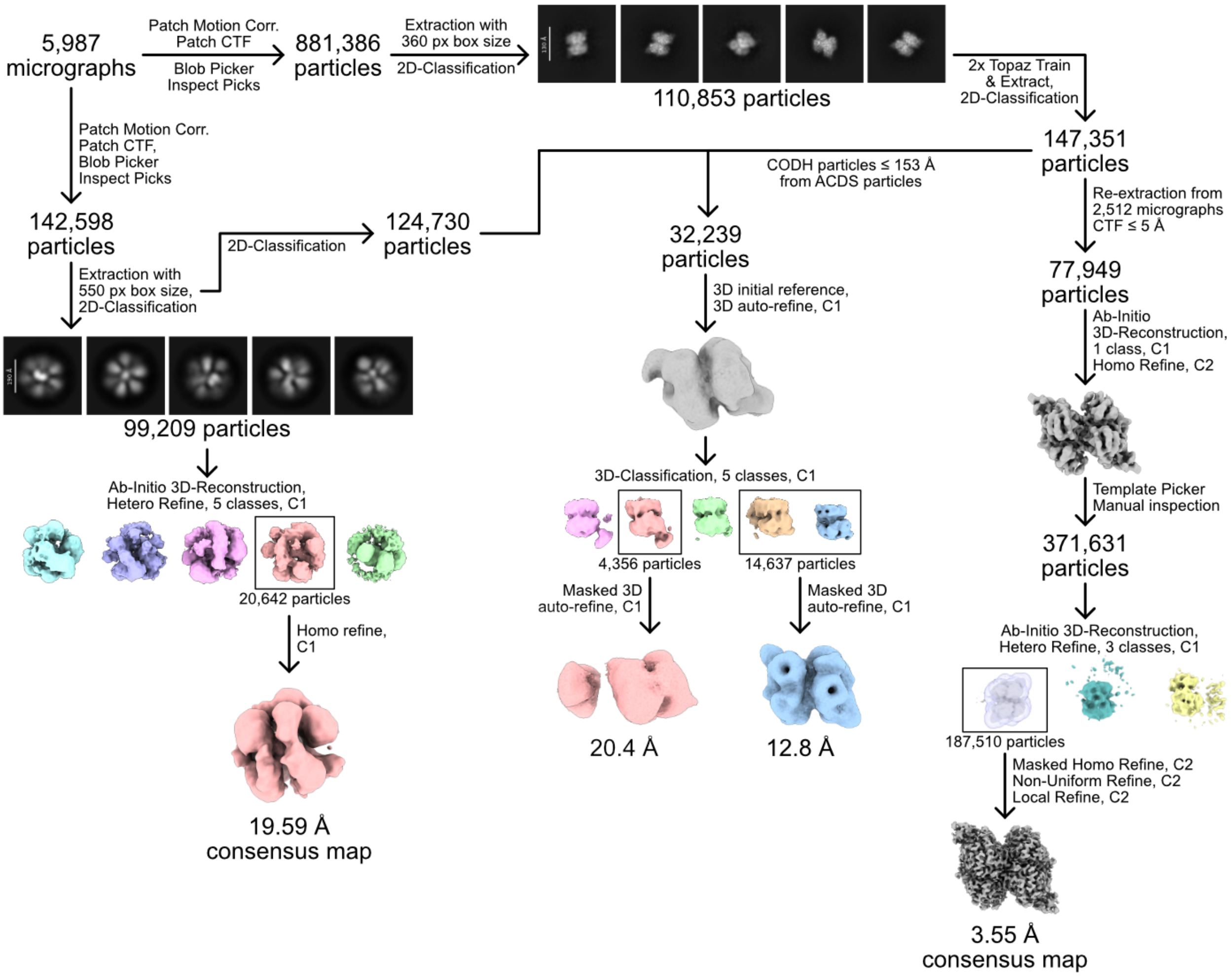

**Extended Data Fig. 3: Cryo-EM processing workflow of CODH and ACDS particles.**

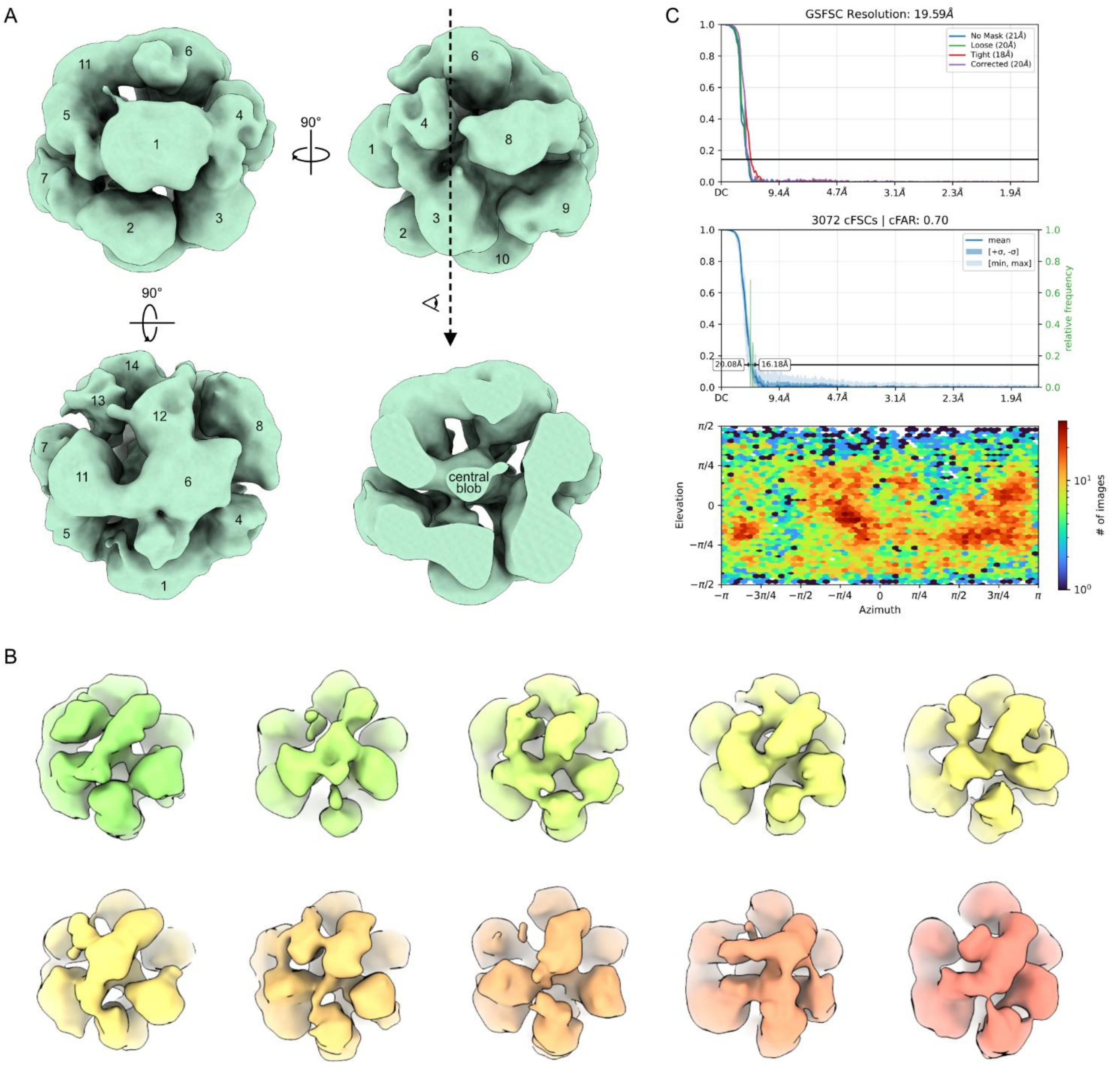

**Extended Data Fig. 4: Cryo-EM consensus map of ACDS particles. (A)** Map view from different orientations, blobs are serially numbered. **(B)** 10-class 3D-classification (Relion) of the 20,642 particles used for the consensus map. **(C)** GS-FSC and orientation distribution plots of the consensus map.

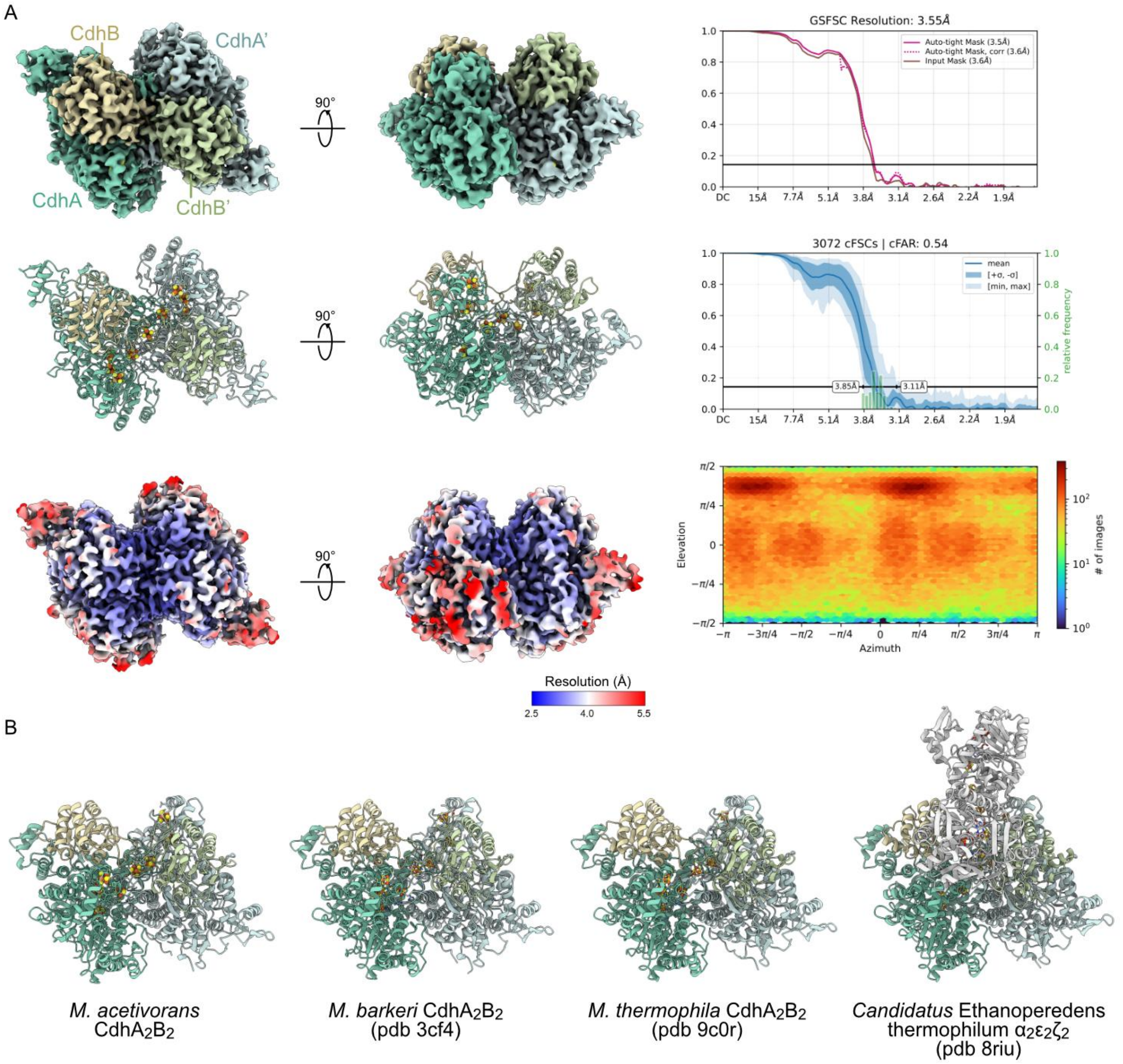

**Extended Data Fig. 5: Cryo-EM structure of M. acetivorans CODH (CdhA2B2). (A)** Map and corresponding model, colored by subunits CdhA, CdhA', CdhB, and CdhB', or colored by local resolution; metal cofactors are shown as spheres. GS-FSC and orientation distribution plots. **(B)** Superposition of the model with published CODH structural models of *M. barkeri* (3cf4, crystal structure), *M. thermophila* (9c0r, EM), and *Candidatus* Ethanoperedens thermophilum (8riu, EM).

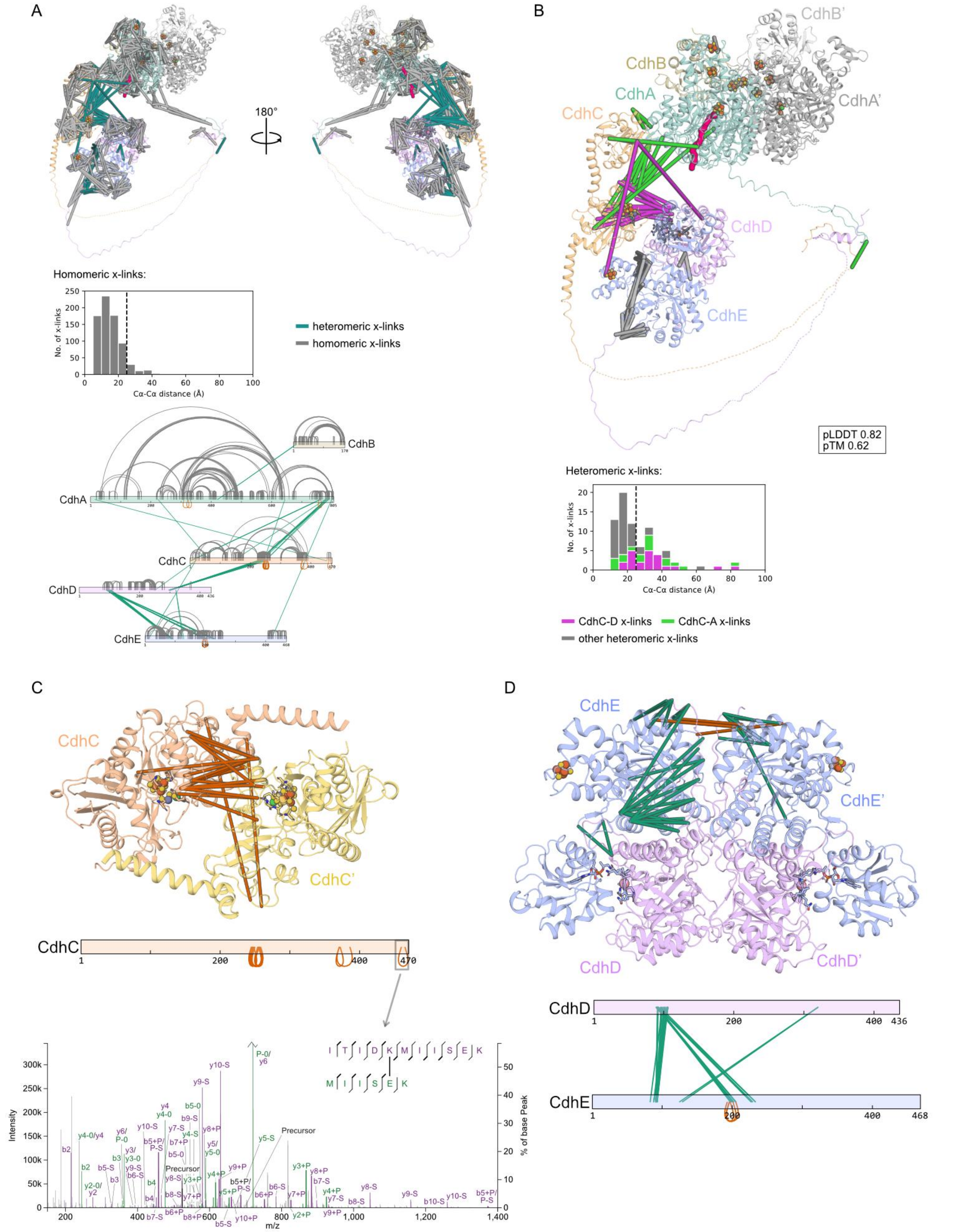

**Extended Data Fig. 6: Crosslinking MS of the ACDS complex. (A)** Plotting of all 777 homomeric non-overlapping (grey) and all 78 heteromeric (green) crosslinks on an AlphaLink2 model of the five ACDS subunits CdhA-E, coloured by subunit (CdhA green, CdhB brown, CdhC yellow, CdhD pink, CdhE blue). Cofactors were created from superpositions of related structures (pdb-ids 1oao and 3cf4 for A- and C-clusters, 2ycl for CdhE corrinoid and [4Fe-4S]) on respective domains. The second CODH CdhAB protomer is shown in grey. The histogram displays the distribution of Cα-

Cα-distances of the homomeric non-overlapping crosslinks in the displayed model; the theoretical distance threshold of 25 Å is indicated by a vertical dashed line. The linear plot displays the locations of all homomeric non-overlapping (grey), overlapping (orange) and heteromeric (green) in the 5 subunits. **(B)** The same CdhABCDE AlphaLink2 model and distance distribution as in (A), but with only heteromeric crosslinks plotted. CdhA-C crosslinks are shown in green, CdhD-C in pink, and all other heteromeric crosslinks in grey. **(C, D)** Dimeric protein models, manually created in ChimeraX, based on homomeric overlapping crosslinks. CA) $CdhC_2$ model of CdhC residues 1-420 with 17 crosslinks (orange) and linear plot displaying the locations of the crosslinks. One of the peptide-spectrum matches underlying the crosslink located in the C-terminal disordered protein region (K460-E465) is displayed. **(D)** $Cdh(DE)_2$ model excluding CdhD residues 1-88, CdhD displayed in pink and CdhE in blue. Heteromeric crosslinks are plotted in green and homomeric crosslinks in orange, a linear plot of the crosslinks' locations is shown. A-cluster, [4Fe-4S] and corrinoid cofactors were created from superpositions of related structures (pdb-ids 1oao for A-clusters, 2ycl for CdhE corrinoid and [4Fe-4S]) on respective domains.

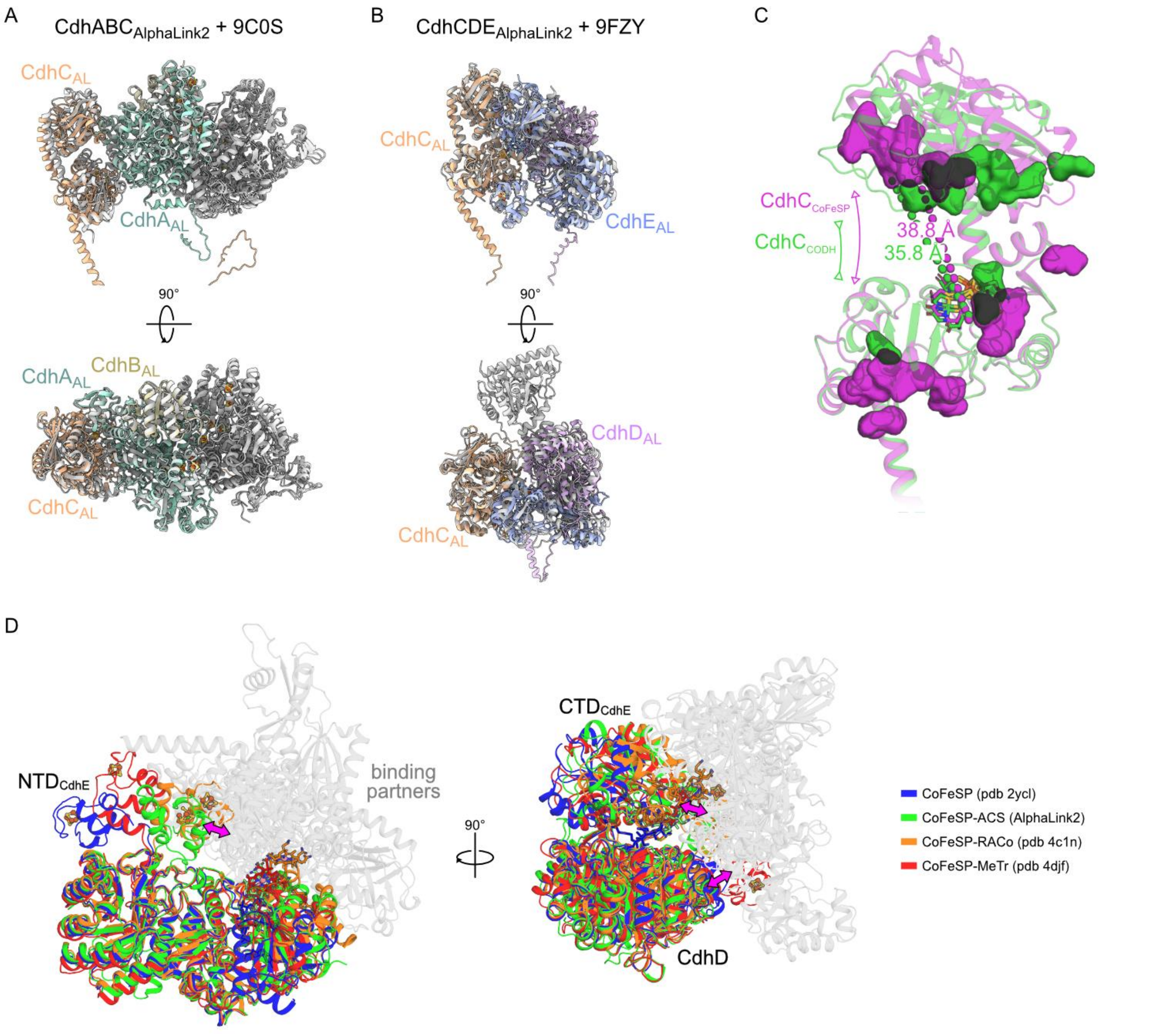

**Extended Data Fig. 7: AlphaLink2 models of ACS-CODH and ACS-CoFeSP interactions resulting from crosslinking MS data. (A)** Superposition of the AlphaLink2 model of CdhC (ACS, yellow) in complex with $CdhA_2B_2$ (CODH, shades of green) with the cryo-EM structure of the ACS-CODH complex from Methanosarcina thermophila (pdb 9c0s, light grey). **(B)** Superposition of the AlphaLink2 model of CdhC (ACS, yellow) in complex with CdhDE (CoFeSP, pink and blue) with the cryo-EM structure of the ACS-CoFeSP complex from Clostridium autoethanogenum (9fzy, light grey). **(C)** Comparison of ACS conformational states in the AlphaLink2 models of ACS in complex with CODH (green) and in complex with CoFeSP (violet); distances of CdhC residue W111 to F195 are shown, highlighting the open (in complex with CoFeSP, violet) and closed (in complex with CODH, green) conformation of ACS; residues involved in the interaction of ACS with CoFeSP (violet) and with CODH (green) in the AlphaLink2 models (within 4 Å distance of the other subunits) are shown as surface representation. **(D)** Superposition of the predicted CdhDE-CdhC (CoFeSP-ACS) AlphaLink2 model (green) with x-ray and cryo-EM structures of CoFeSP alone (pdb 2ycl, blue), CoFeSP in complex with its reductive activator (RACo, pdb 4c1n, orange), and CoFeSP in complex with a methyl transferase (MeTr, pdb 4djf, red). Commonly found protein-protein interfaces are denoted with pink arrows, cofactors (from structures or superposition with pdb 2ycl) are shown as spheres and sticks.

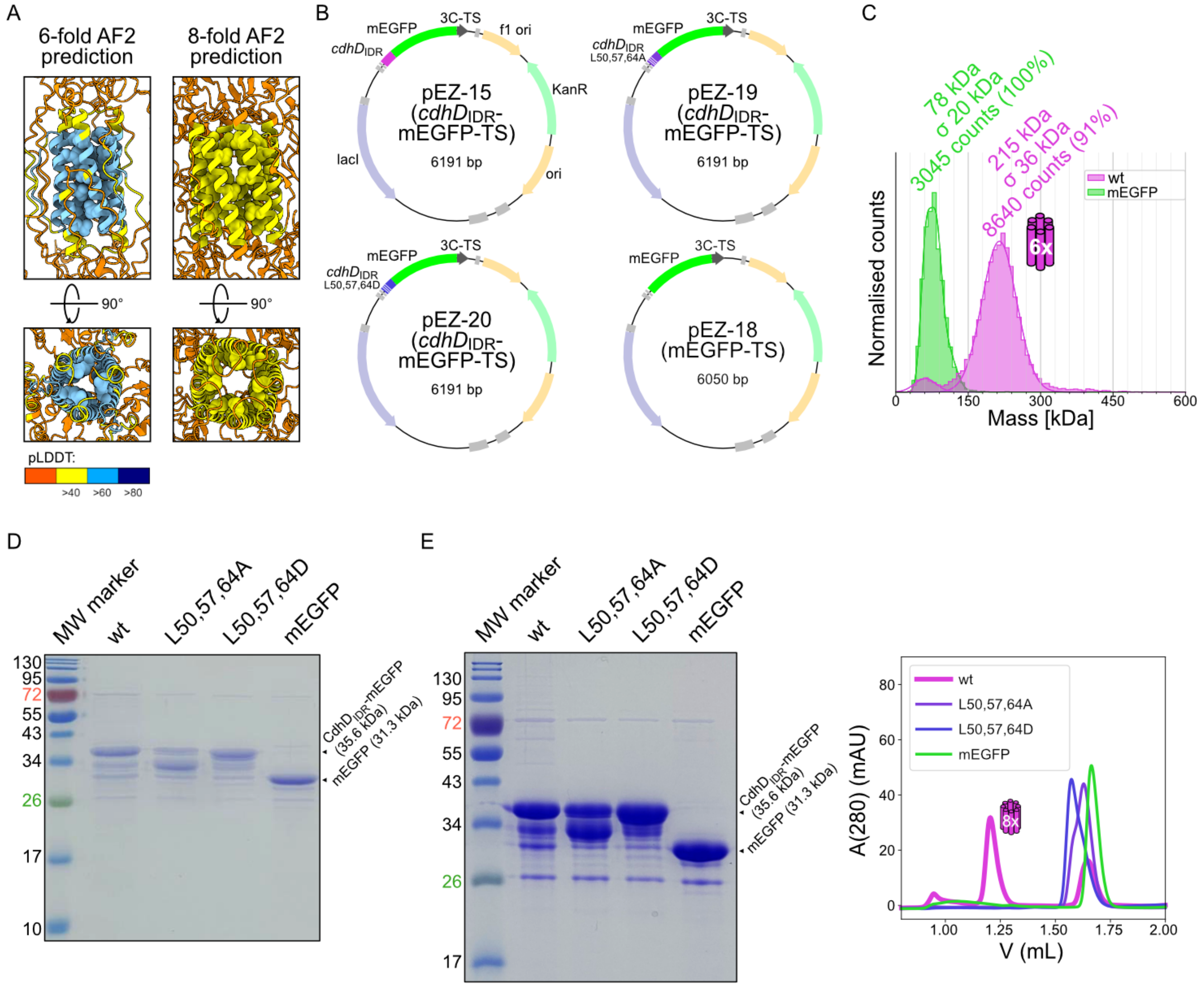

**Extended Data Fig. 8: Extended data on experiments with the N-terminal disordered region (IDRs) of ACDS subunit CdhD. (A)** AlphaFold2 predictions of 6- and 8-fold stoichiometry of the three ACDS terminal disordered regions (CdhA residues 1-47, CdhC residues 401-470, CdhD residues 1-112), colored by pLDDT score; CdhD Leu residues 50, 57 and 64 are depicted as spheres. **(B)** Plasmid maps for expression of $\mathrm{CdhD_{IDR}}$-mEGFP-TS fusion constructs, of CdhD residues 43-88 fused to mEGFP; from left to right, fusion of the wildtype (wt) CdhD sequence, Leu residues 50, 57 and 64 mutated to Ala, mutated to Asp, and mEGFP without CdhD-fusion. **(C)** Mass photometry histogram of purified wt $\mathrm{CdhD_{IDR}}$-mEGFP-TS (pink) and mEGFP-TS (green); Fitted mass photometry peaks are labelled with their mean ± standard deviation molecular weights (MWs); The 215 kDa peak corresponds to hexameric $\mathrm{CdhD_{IDR}}$-mEGFP-TS (theoretical MW 214.8). **(D)** Coomassie-stained SDS-PAGE of the four purified fusion constructs, replicate 1, and **(E)** SDS-PAGE and S200 Increase 3.2/300 gel filtration chromatogram of replicate 2; The multiple bands observed in SDS-PAGE are either a product of partial degradation of unstructured protein regions, or of incomplete denaturation of mEGFP, since green fluorescent protein bands were observed even after heat denaturation for 15 min in SDS-containing buffer.

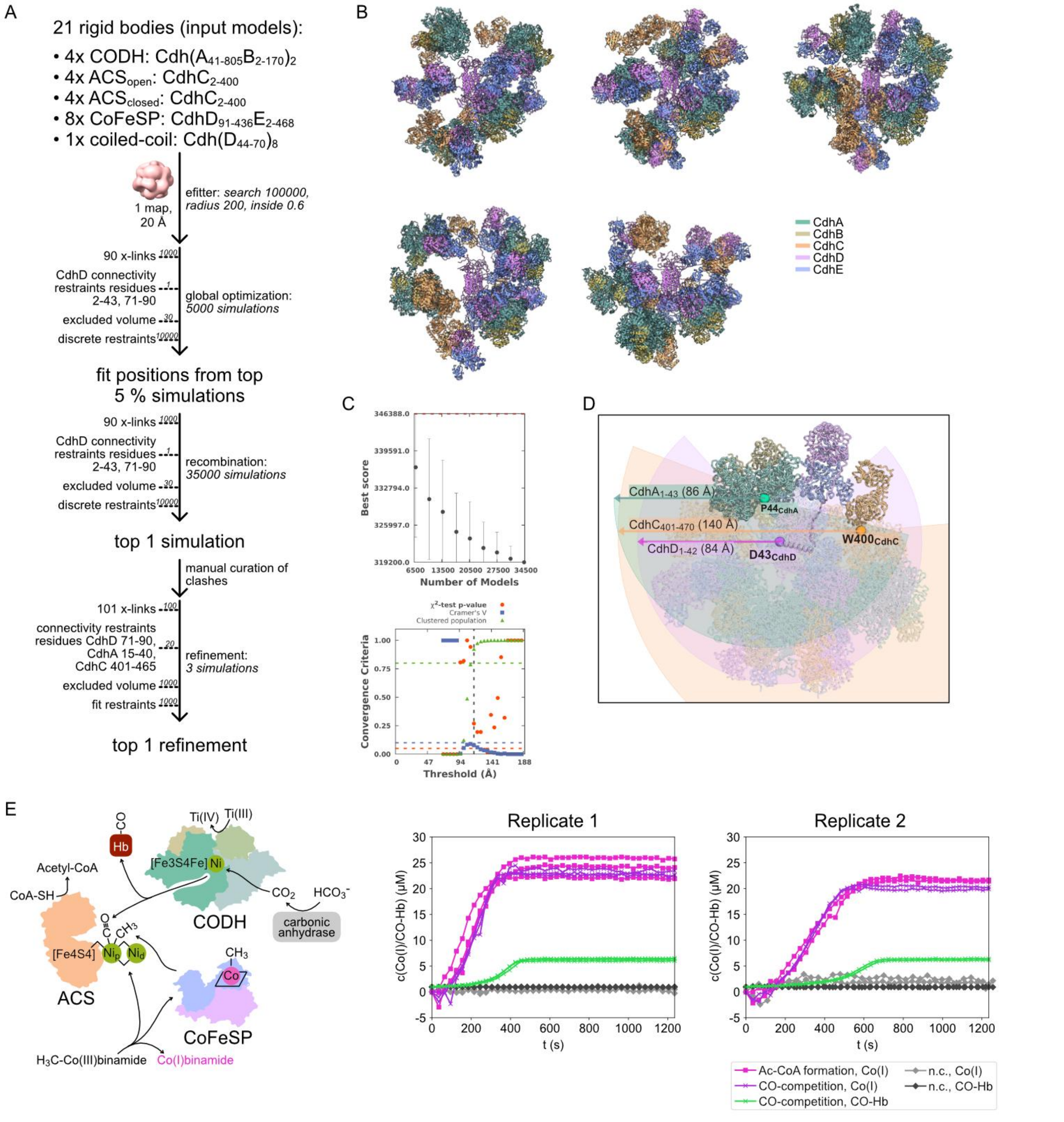

**Extended Data Fig. 9: Assembline integrative modelling of the ACDS complex to explain the efficient CO and methyl-transfer between subcomplexes. (A)** Modelling workflow; Numbers above the dashed lines denote weights of the different restraints in the scoring function. **(B)** Top 5 refined models, coloured by subunit. **(C)** Sampling convergence and precision plots after the recombination simulations. **(D)** Depiction of areas theoretically accessible by disordered regions of subunits CdhD (M1 to D43, the protein region N-terminal of the coiled coil, pink), CdhA (M1 to P44, the N-terminal region, green), and CdhC (W400 to K470, the C-terminal region, yellow), assuming a relaxed backbone conformation with 2.0 Å per residue. The relevant residues are highlighted as colored spheres. **(E)** Schematic representation of the assessed catalyzed reactions, and replicates of the assay; the acetyl-CoA formation assay contained 5 mM NaHCO3, 0.3 mg/L carbonic anhydrase, 5 mM Ti(III)-EDTA, 200 µM CoA, 50 µM methyl-Co(III)binamide and 0.5 µM ACDS; the CO competition assay additionally contained 7 µM hemoglobin. Abbreviations: (carboxy)hemoglobin ((CO-)Hb); acetyl-coenzyme A (Ac-CoA); negative control (n.c.); acetyl-CoA synthase (ACS); CO dehydrogenase (CODH); corrinoid iron-sulfur protein (CoFeSP).

**Extended Data Table 1: Cryo-EM data collection, refinement and validation statistics**

| | **CODH ($CdhA_2B_2$)** (EMD-58601) (PDB: 31OX) | **ACDS ($CdhA_8B_8C_8D_8E_8$)** (EMD-58600) |
|---|---|---|
| **Data collection and processing** | | |
| Magnification | 60,000 | 60,000 |
| Voltage (kV) | 200 | 200 |
| Electron exposure ($e^-/Å^2$) | 50 | 50 |
| Defocus range (μm) | 0.5-2.5 | 0.5-2.5 |
| Pixel size (Å/px) | 0.85 | 0.85 |
| Symmetry imposed | C2 | C1 |
| Initial particle images (no.) | 881,386 | 142,598 |
| Final particle images (no.) | 187,510 | 20,642 |
| Map resolution (Å) | 3.55 | 19.59 |
| FSC threshold | 0.143 | 0.143 |
| Map resolution range (Å) | 3.0-5.5 | - |
| **Refinement** | | |
| Initial model used | AlphaFold2 | Integrative |
| Model resolution (Å) | 3.3 | 44.4 |
| FSC threshold | 0.143 | 0.143 |
| Model resolution range (Å) | 2.8-3.7 | - |
| Map-model CC (box) | 0.73 | 0.62 |
| Map sharpening *B* factor ($Å^2$) | 116.2 | - |
| Model composition (no.) | | |
| Non-hydrogen atoms | 14,212 | 141,276 |
| Protein residues | 1,848 | 18,272 |
| Ligands | 9 | 76 |
| *B* factors ($Å^2$) | | |
| Protein | 24.18 | 56.12 |
| Ligands | 24.34 | 30.27 |
| R.m.s. deviations | | |
| Bond lengths (Å) | 0.002 | 0.017 |
| Bond angles (°) | 0.669 | 1.494 |
| Validation | | |
| MolProbity score | 1.38 | 1.99 |
| Clashscore | 4.23 | 6.75 |
| Rotamer outliers (%) | 0.53 | 1.43 |
| Ramachandran plot (%) | | |
| Favored | 97.00 | 91.58 |
| Allowed | 3.00 | 6.92 |
| Disallowed | 0.00 | 1.51 |
